# An integrated multi-omic analysis of iPSC-derived motor neurons from C9ORF72 ALS patients

**DOI:** 10.1101/2020.11.01.362269

**Authors:** The NeuroLINCS Consortium, Loren Ornelas, Emilda Gomez, Lindsay Panther, Aaron Frank, Susan Lei, Berhan Mandefro, Maria G Banuelos, Brandon Shelley, Julia A Kaye, Leandro Lima, Stacia Wyman, Ryan G Lim, Jie Wu, Jennifer Stocksdale, Malcolm Casale, Victoria Dardov, Andrea Matlock, Vidya Venkatraman, Ronald Holewenski, Pamela Milani, Miriam Adam, Brook T Wassie, Andrew Cheng, Alyssa N Coyne, J. Gavin Daigle, Johnathan Li, Stephanie Yang, Veerle Cox, Mark Wilhelm, Thomas E Lloyd, Lindsey Hayes, Jacqueline Pham, Renan Escalante-Chong, Alex Lenail, Karen Sachs, Natasha Leanna Patel-Murray, Divya Ramamoorthy, Terri G Thompson, NYGC ALS Consortium, Steven Finkbeiner, Ernest Fraenkel, Jeffrey D Rothstein, Druv Sareen, Jennifer E Van Eyk, Clive N Svendsen, Leslie M. Thompson

## Abstract

Neurodegenerative diseases present a challenge for systems biology, due to the lack of reliable animal models and the difficulties in obtaining samples from patients at early stages of disease, when interventions might be most effective. Studying induced pluripotent stem cell (iPSC)-derived neurons could overcome these challenges and dramatically accelerate and broaden therapeutic strategies. Here we undertook a network-based multi-omic characterization of iPSC-derived motor neurons from ALS patients carrying genetically dominant hexanucleotide expansions in *C9orf72* to gain a deeper understanding of the relationship between DNA, RNA, epigenetics and protein in the same pool of tissue. ALS motor neurons showed the expected *C9orf72*-related alterations to specific nucleoporins and production of dipeptide repeats. RNA-seq, ATAC-seq and data-independent acquisition mass-spectrometry (DIA-MS) proteomics were then performed on the same motor neuron cultures. Using integrative computational methods that combined all of the omics, we discovered a number of novel dysregulated pathways including biological adhesion and extracellular matrix organization and disruption in other expected pathways such as RNA splicing and nuclear transport. We tested the relevance of these pathways *in vivo* in a *C9orf72* Drosophila model, analyzing the data to determine which pathways were causing disease phenotypes and which were compensatory. We also confirmed that some pathways are altered in late-stage neurodegeneration by analyzing human postmortem C9 cervical spine data. To validate that these key pathways were integral to the C9 signature, we prepared a separate set of *C9orf72* and control motor neuron cultures using a different differentiation protocol and applied the same methods. As expected, there were major overall differences between the differentiation protocols, especially at the level of in individual omics data. However, a number of the core dysregulated pathways remained significant using the integrated multiomic analysis. This new method of analyzing patient specific neural cultures allows the generation of disease-related hypotheses with a small number of patient lines which can be tested in larger cohorts of patients.

## Introduction

Modeling neurological diseases using induced pluripotent stem cell (iPSC) technology offers a unique platform to study the process of pathogenesis. Rather than using artificially expressed human disease genes in mice or end stage post-mortem tissues from patients, the generation of new neurons and astrocytes from patient-specific cells allows for discovery of the earliest genesis of disease signatures. One neurodegenerative disease group that has been modeled extensively using iPSCs is the motor neuron disorders. Adult onset motor neuron diseases include amyotrophic lateral sclerosis (ALS), where motor neurons degenerate late in life, inevitably leading to paralysis and asphyxiation. Genetic underpinnings have been identified in ∼15% of ALS cases^1^. Of these, the most common mutation is a hexanucleotide repeat expansion (HRE) in the first intronic region of *C9orf72,* which accounts for over 40% of all known familial and 10% of known sporadic forms of the disease. In healthy individuals, fewer than 24 copies of the GGGGCC HRE are present within the first intron of the *C9orf72* gene. However, in disease this GGGGCC sequence is expanded hundreds to thousands of times. While there is no known correlation between HRE length and disease severity, this intronic expansion leads to three pathologic hallmarks of C9orf72 ALS/FTD. First, the HRE has been shown to lead to reduced C9orf72 RNA and protein expression, leading to a loss of function of the C9orf72 gene. Second, there is a gain of toxic function via the bidirectional transcription of the GGGGCC HRE leading to the production of toxic G_4_C_2_ and G_2_C_4_ repeat RNA species which are thought to sequester and impair the function of RNA binding proteins. Third, there is a gain of toxic function via the non-canonical RAN translation of repeat RNAs to produce 5 toxic dipeptide repeat proteins (Poly(GR), Poly(GA), Poly(GP), Poly(PR), and Poly(PA)) which are proposed to impair multiple cellular processes^2–4^. While much is known about the mutation and abnormal proteins that are produced by its transcripts^5^, it still remains unclear as to how repeats in *C9orf72* lead ultimately to neuronal dysfunction and death.

Some of the first disease modeling studies showed that iPSCs could be generated from early onset motor neuron diseases such as spinal muscular atrophy and that these motor neurons exhibited disease-specific cell death *in vitro*^6–11^. Interestingly, for later onset motor neuron diseases such as ALS, initial studies of iPSC models did not show any overt death in motor neurons^12^. However, for inherited forms of ALS such as *C9orf72* repeat expansions (C9) there were specific changes in neuronal activity, gene expression, and cellular processes^13–18^. More recently, stressors, such as trophic factor withdrawal, have led to cell death phenotypes^17^, although it is not clear how these stressors relate to human disease onset and progression. Subsets of sporadic ALS patients also showed phenotypic changes including reduced fiber outgrowth at later time points in culture^19^, although a comprehensive omics analysis was not performed and *C9orf72* cases were not included. These iPSC models provide a unique opportunity to examine the *molecular* changes that occur due to ALS-causing genes in motor neurons. Whereas post-mortem studies^20–24^ have provided important insights into these processes, patient samples often represent a late stage of the disease with extensive degeneration, which may not exhibit molecular or cellular signatures directly associated with the initiating events that cause the disease. By contrast, cells derived from iPSCs can provide insights into the earliest stages of neurodegeneration, opening a window into the period when therapeutics might have the greatest benefit.

Although iPSCs represent critical models that represent disease within their human context, it is clear that one of the major challenges for the field is patient to patient variation in iPSC lines and major challenges regarding the reliable and reproducible differentiation of motor neurons from iPSC’s. Batch to batch variation in differentiations are driven by hard to control factors such as small variances in the multitude of small molecules and other media components along with plating densities and technical variations in feeding and handling. The combined heterogeneity in iPSC state and differentiation protocols means even within the same lab it is difficult to control completely, and that comparing data between labs using different protocols is almost impossible.

The goal of the current study was to use a multi-omic approach to test whether a network-based analysis would facilitate identification of early pathogenic events in C9 ALS where the signal was strong enough to rise above the noise of the system. We differentiated cells for all the assays at once, and then divided the cells for each omics, using stringent quality control measures in both experimental and analytical steps. We also developed an integrative approach that combines multi-omic data using network-based algorithms. Significant signals emerged even with a small sample size that were validated in human postmortem cervical spine transcriptomes. We then carried out experiments in a Drosophila model to test whether the “hits’ were relevant to C9 ALS and if they were pathological changes or compensatory responses to neurodegeneration, and we integrated these data into a map of key pathways changes in C9 patient cells.

A critical test of this approach is whether similar findings can be found in cells from different donors or differentiation states. To that end, we differentiated a different set of lines using a different differentiation protocol – but applied the same integrative approach. Despite the many experimental differences, our computational approach confirmed a set of causal and compensatory changes that were detected in both sets of samples. These studies support the feasibility of a network-based multi-omic approach to generate disease related hypotheses and to establish the paradigm. More patients will now be required to validate these pathways further in continuing experiments. As part of the NIH-funded NeuroLINCS consortium, all of the data sets along with the data integration have been posted to a portal for data sharing of this unique resource http://neurolincs.org and the iPSC lines are all available at https://stemcells.nindsgenetics.org/. This study has led to the formation of Answer ALS where1000 iPSC lines are currently being generated and analyzed using similar integrative analyses in familial and sporadic ALS patients to explore the power of this approach further, stratify ALS subpopulations and identify therapeutic targets.

## Results

### Generation and characterization of iPSC lines

An initial set of four C9-ALS lines and three control iPSC lines generated from patient fibroblasts and reported on previously ^15^ were used for the majority of this study. We have previously shown that motor neurons derived from these C9-ALS lines showed RNA foci, physiological changes that including a diminished capacity to fire continuous spikes and changes in specific genes based on an RNAseq experiment^15^. We used this data as a baseline for the extended studies in the current paper using a much wider range of omics analysis and network-based analysis. In addition, we extended the number of lines to an additional 7 controls and 6 C9-ALS iPSC lines but using PBMC’s as the starting cell source for a replication cohort. All lines were generated using episomal plasmid-based reprogramming methods. Lines (list in **Supplementary Fig. 4a and b**) were differentiated to motor iMNs as described in methods, with no overall differences in cell markers (**Figure 1a,b,c**). The lines retained their repeat expansion mutation following reprogramming as described previously (^15^, https://www.biorxiv.org/content/10.1101/2020.04.27.064584v1). All iPSC lines maintained normal karyotypes as determined by G-band karyotyping (**Supplementary Fig. 4a and b)** and the identity of iPSCs and differentiated iMNs were confirmed to match the parent fibroblasts or isolated PBMCs by DNA fingerprinting (**Supplementary Fig. 5a and b**).

**Figure 1:**
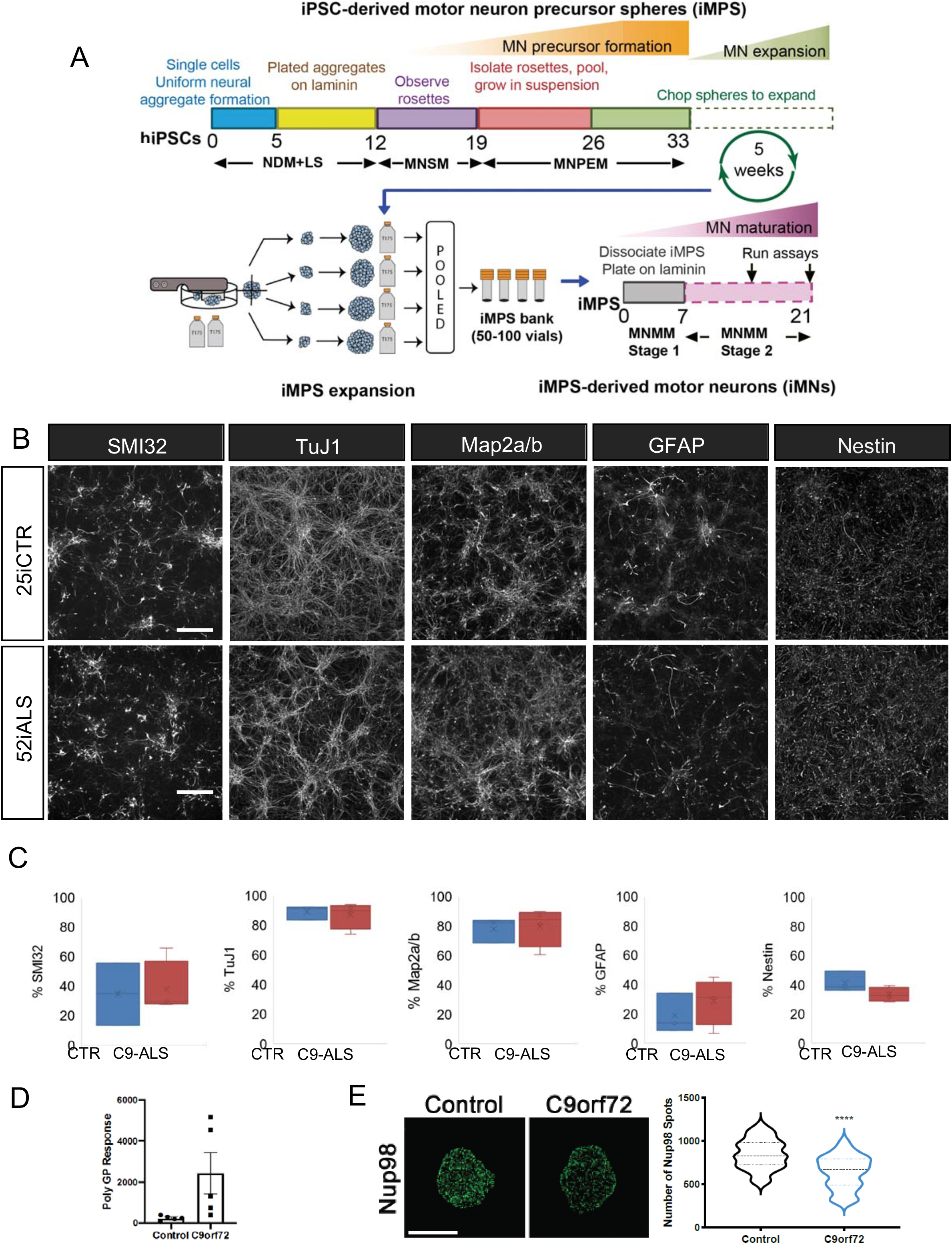
iPSC differentiations. (A) Schematic of protocol for iPSC differentiation into motor neuron cultures used by NeuroLINCS for transcriptomics, proteomics and ATAC-seq assays. The iPSC-derived motor neuron precursor spheres (iMPS) were dissociated into single cells from C9-ALS and healthy patient iPSC lines and plated on laminin substrate to differentiate further into motor neuron (iMN) cultures over 21 days. (B) Representative images of iMNs from control (25iCTR) and C9-ALS (52iALS) iMPS shows consistent distribution of neural cell populations marked by SMI32, TuJ1, Map2a/b, GFAP and nestin. Scale bars are 50 µm. (C) Box plots quantifying levels of SMI32, TUBB3 (Tuj1), GFAP, nestin and Map2a/b in control and C9-ALS iMN cultures from the individual iPSC lines. Two-sided unpaired t test with Welch’s correction (CTR n=3 and C9-ALS n=4).. (D) Poly(GP) DPR levels as determined by MSD ELISA assay in iMNs (from CS0188,CS0594,CS0702,CS29,CS52,CS7VCZ). (E) Maximum intensity projections from SIM imaging of Nup98 in nuclei isolated from control and C9orf72 iMNs. Quantification of Nup98 spots. N = 3 control and 3 C9orf72 iPSC lines, 20 NeuN+ nuclei/line. Student’s t-test was used to calculate statistical significance. Scale bar = 5 um.

We also evaluated C9 phenotypic signatures to determine that the iMNs produced relevant C9orf72 pathology. We first evaluated dipeptide repeat species (DPR) species using immunoassays to evaluate the expression of Poly(GP)^25^. Compared to controls, C9orf72 iMNs produce significantly more Poly(GP) (**Figure 1d**). In accordance with previous reports, the levels of Poly(GP) vary among individuals. Impaired nucleocytoplasmic transport and alterations in the expression and localization of specific nucleoporins which make up the nuclear pore complex to govern functional nucleocytoplasmic transport have emerged as a prominent pathologic hallmark of multiple neurodegenerative diseases including C9orf72 ALS/FTD. To verify that nucleoporin components are altered, we performed super resolution structured illumination microscopy (SIM) on nuclei isolated from control and C9orf72 iMNs immunostained for Nup98 as previously described^26^. Nuclear preparations and SIM are required to identify changes in nuclear pore proteins; these are not readily observed through proteomic analysis. In comparison controls, we observed a significant reduction in the nuclear expression and localization of Nup98 in C9orf72 iMNs (**Figure 1e**) similar to previous pathological observations in iPSNs and postmortem tissue^26^.

### Whole genome sequencing shows no known disease-associated ALS variants

WGS was performed on the initial set of C9-ALS and control iPSC lines to establish methodologies and provide a reporting of variants in disease–modifying genes to help elucidate and interpret line to line variability, despite the fact that the vast majority of these variants are benign or of unknown significance. A novel computational pipeline was used to annotate the variants in the genomes of the control and C9-ALS lines relative to reference human genomes (see WGS methods). The number of single nucleotide polymorphisms (SNPs) was within the expected range, and there were no overt genetic abnormalities. Across all lines, we found 11,260,464 variants with 9,197,462 variants in the control lines and 8,818,235 variants in the C9-ALS lines. Thus, there was an average of 5.4 million variants per line, which is consistent with the variation that has been previously observed in human genomes^27^. After applying annotation (see methods), we filtered for exonic functional variation (Table 1). There were 57,910 exonic functional variants in the controls, and 12,898 were rare (less than 1%) or novel (no frequency information). There were 55,815 exonic functional variants in the C9-ALS lines, and 8,225 were rare or novel (**Supplementary Table 6**). Next, we investigated if any of the lines had genetic variants previously associated with ALS, and we found 3 variants in OPTN, ALS2 and DIAPH3 that have been associated with ALS, but are also found at relatively high frequency (>2%) in the general population (**Supplementary Table 7**). Other variants in ALS-associated genes were observed, but none that were known previously to be disease-associated or causing. However, of interest, the 52i ALS line contains the APOE-ε4 allele (rs429358) (C130R) which is associated with an increased risk of developing Alzheimer’s disease^28^. We next applied the American College of Medical Genetics gene criteria to identify likely pathogenic (LP) or Pathogenic (P) variants (**Supplementary Table 8**). Although a subset of these variants is in ALS genes that are listed in the ASLoD database^29^, to our knowledge none of these variants are expected to confer risk of developing ALS. WGS analysis of the patient cell lines revealed no pathogenic or likely pathogenic variants hence there is no indication that these variants confer risk or influence risk of developing ALS.

### Transcriptomic analysis reveals known and novel pathways related to C9

To identify the earliest molecular changes in the differentiated ALS iMNs, we carried out parallel multi-omics analyses. RNA sequencing revealed specific transcriptomic signatures associated with the C9 lines (**Supplementary Table 9**). Total RNASeq (Ribo-Zero rRNA depletion) was carried out on the distributed iMN pellets as described in methods. Statistical analysis of differential expression was analyzed using DEseq2. We found 828 differentially expressed transcripts (271 downregulated and 557 upregulated) between C9-ALS and control iMNs (FDR<0.1), of which 704 were annotated as protein-coding in Uniprot (**Supplementary Table 9 & Supplementary Fig. 6a,b)**. Of these 828 DEGs, C9orf72 was not differentially expressed. Exploratory analysis of gene expression levels was carried out using Hierarchical clustering (**Figure 2a**). To begin to understand the effect of the C9 mutation on a multicellular culture, genes that were significantly different between C9 and control samples were used for Cell Type-Specific Expression Analysis (CSEA^30^) (**Supplementary Fig. 6c**). CSEA revealed an enrichment of cortical and motor neuron specific gene expression. Next, gene ontology (GO) analysis was conducted to determine the functional role of these genes, using GOrilla on the 704 DEGs, revealing an enrichment in Extracellular Matrix (ECM) and cell adhesion terms which included: ECM disassembly, ECM organization, collagen binding, and focal adhesion (**Figure 2c; Supplementary Fig.6d,e**). A previous study^15^ using the same C9 iPSC-derived motor neurons and 2 of the same controls, but a very different differentiation protocol, also showed dysregulation; in this case 66 genes between 4 ALS and 4 control samples with a fold change of >2 and p-value <0.05 were dysregulated. Of those 66 genes, 8 genes overlapped with the 828 DEGs from our study, although in different directions. While specific genes differed, even with this small number of overlapping genes from different studies using distinct differentiation protocols and different RNASeq platforms, GO enrichment analysis revealed an enrichment for extracellular region in the 66 genes, similar to our analysis suggesting that at the individual gene level, batch, differentiation and study effects make direct comparisons difficult, but, at the global pathway level, disease specific signatures remain. This formed the rational for extending these large network analysis further to include proteomics and ATAC seq with the hypothesis that more data would provide consistent signatures that separated ALS from control and would enable discovery of mechanistic disease relevant pathways.

**Figure 2:**
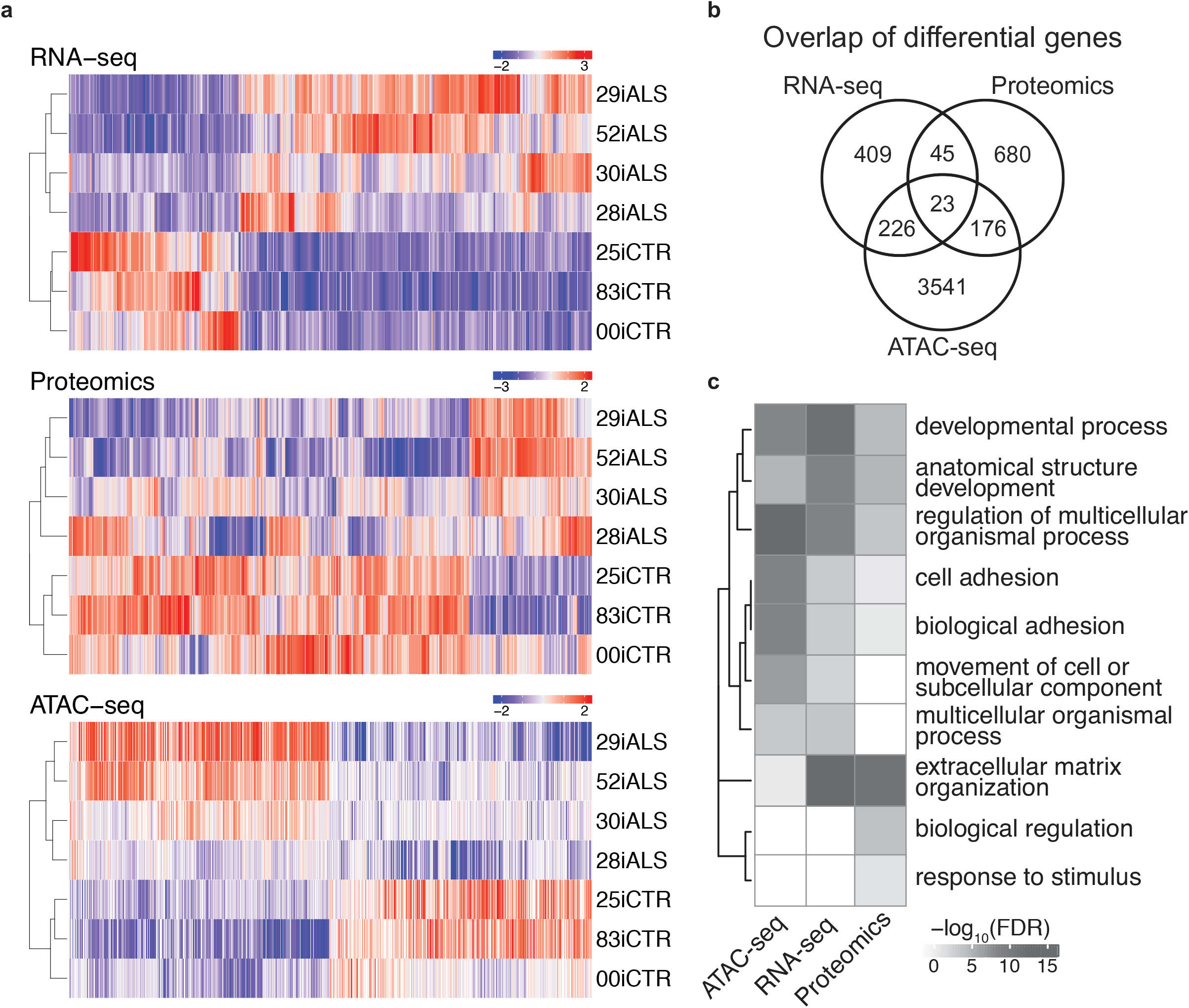
OMIC assays. (A) Hierarchical clustering of RNA-Seq, Proteomics, and ATAC-seq signals normalized by z-score. (B) Venn diagram of differential genes or proteins from each assay. Each differential ATAC-seq peak was assigned the nearest protein coding gene (up to a limit of 50kb from the TSS). (C) Top GO term enrichments for each assay reveal common biological processes.

To identify potential regulators controlling the differential expression of these ECM related genes, Ingenuity pathway analysis (IPA) upstream regulator analysis was conducted. Some of the top predicted regulators identified include SMADs (transforming growth factor beta (TGFβ) signaling), mitogen-activated protein kinase 1 (ERK), and nuclear factor kappa B (NFκB). Network-based analysis of upstream regulators and gene targets showed a TGFβ, AP-1 transcription factor subunit (AP1), erb-b2 receptor tyrosine kinase 2 (ERBB2), plasminogen activator, urokinase receptor (PLAUR), and neuregulin 1 (NRG1) network that were again predicted to regulate many of the ECM and cell adhesion related DEGs (**Supplementary Fig. 6d**). Notably, NRG1 was identified as a major hub gene that could regulate other upstream regulators and directly regulate *ACTIN* and *INTEGRIN* expression, each of which was upregulated in the ALS iMNs. Matrix metalloproteinases (*MMP*s) were significantly dysregulated, in all cases showing increased expression, and were downstream of the NRG1 hub (**Supplementary Fig. 6e**). We further investigated dysregulation of these *MMPs* and found that their corresponding substrates (*e.g.*, *LAMININs*, *COLLAGENs*) were also upregulated (**Supplementary Fig. 6e**). These data indicate a potential role for NRG1 in the dysregulation of ECM and cell adhesion-related genes in ALS iMNs, as suggested previously in mouse models of ALS^31^.

The use of total RNASeq allowed further analysis of the transcriptomic data focused on alternative splicing. This analysis was conducted using MATS^32^. **Supplementary Fig. 6f** shows the percentage of significant alternative splicing events found in the ALS iMNs compared to controls, showing a high percentage of Exon skipping (ES, 57%) and Intron retention (RI, 26%). This same pattern was previously identified as enriched in studies using human familial ALS and sporadic ALS patient tissue^23^. GO enrichment analysis of these alt spliced genes revealed some similar terms found in the gene level differential analysis like cell adhesion, but also unique terms related to RNA processing, axonal guidance, and translation (**Supplementary Figs. 6g-i**).

### Proteomics shows ECM and mRNA processing dominate protein changes

A sample specific library using DDA based acquisition files was compiled and DIA-MS samples were run against the peptide library. Data quality was assessed by MS1 and MS2 total ion current, normalized protein intensity distribution, number of unique and shared hits identified, and correlation between ALS and control lines (**Supplementary Fig. 7a-d**). Using this method, we were able to identify 3844 unambiguous proteins based on 23,436 unique peptides (**Supplementary Fig. 7a**). MAP DIA software was then used to determine relative peptide and protein amounts within the samples, as well as log2FC between C9 ALS and control using transition level data. Using a 1% FDR, 95% confidence interval and 0.6 abs(log2FC) cutoff, a final list of 924 differentially expressed proteins was obtained; C9orf72 was not differentially expressed. Hierarchical clustering of differential protein intensity values showed similar groupings between biological replicates for ALS and Control samples as seen for RNA-seq and ATAC-seq (**Figure 2a**). Interestingly, unbiased analysis of all measured proteins resulted in separation between control and ALS groups (**Supplementary Fig. 7f**).

A small subset of the differentially expressed proteins (6.8%) had overlap with both the ATAC-seq and RNA-Seq differentially expressed genes (**Fig. 2b**), specifically 68 common differentially expressed genes/proteins (45 between RNA and protein and 23 between all omics data sets) (**Fig. 7c**). The fact that mRNA levels can be poor predictors of protein abundance has been documented previously (e.g.^33^ and references therein), further highlighting the importance of incorporating integrated analytical approaches. The fold change values of these overlapping terms have a correlation R^2^ = 0.76, suggesting that most of the differentially expressed terms that are common have concordant fold change values and directionality (**Supplementary Fig.7e**). Of these common proteins, downregulated proteins (13) did not yield any GO enrichment terms (**Supplementary Table 10b**). Common upregulated proteins/genes (55) show enrichment in extracellular matrix terms (**Supplementary Table 10a**).

The role of the extracellular matrix is further supported by the analysis of the 856 differentially expressed proteins that did not overlap with differentially expressed genes. Of these, 183 proteins were upregulated and enriched for extracellular matrix proteins (**Supplementary Table 10c**), similar to the transcriptomic analysis. In addition, network-based analysis of all differentially expressed proteins (924) by IPA revealed predicted upstream regulators, including TGFβ and SMAD4 (**Supplementary Table 10e**), which in turn regulate many of the extracellular matrix proteins identified in the differential protein analysis and integrated –omics described below.

The remaining unique subset of proteins (674 differentially expressed proteins) were downregulated and showed enrichment for poly(A) RNA binding, RNA binding, RNA and mRNA splicing (**Supplementary Table 10d**). Additionally, IPA analysis of the differential proteins (924) shows predicted inhibition of RNA/mRNA splicing based on downregulation of proteins associated with this pathway (**Supplementary Fig. 7g, Supplementary Table S10f-g**). Lastly, proteins associated with alternative splicing of mRNA are dysregulated, with most of these proteins decreasing in ALS. Taken together, this could imply that these downregulated proteins are associated with the altered exon usage and alternative splicing in ALS found in the transcriptomic analysis.

### Epigenetic changes due to C9 expression seen with ATAC-seq

We sought to study the accessible chromatin landscape in C9 patients and controls. The density of transposase Tn5 cleavage fragments provides a continuous measurement of chromatin accessibility via ATAC-seq (**Supplementary Fig. 8a**). Analysis of the open chromatin data identified 128,299 peaks that were active in 2 or more ALS or control samples. Approximately 14% (18,407) accessible regions localize to gene promoters as defined by GENCODE^34^; 27% (34,543) lie within 2.5 kb of a TSS. Nearly half of the peaks lie in intronic regions, while about a third lie between genes (**Supplementary Fig. 8b**).

To study alterations in chromatin accessibility in the disease state, we identified and characterized peaks with significantly changed accessibility between C9 and control samples. Roughly 12% (15,814 peaks; FDR<0.1) of all peaks were found to be differentially open, of which approximately one half (7,937) were less accessible in C9 samples (**Supplementary Fig. 8c**). Hierarchical clustering of differentially open regions revealed similar groupings of patient samples as in RNASeq and proteomics (**Figure 2a**). Correlation coefficients were 0.46 for RNA and ATAC and 0.13 for protein and ATAC, with both comparisons indicating same direction. Differentially accessible peaks were biased away from regions near TSSs, with only 5.0% (783) annotated to promoters (**Supplementary Fig. 8c**). Examples of changing chromatin accessibility in ALS versus control lines can be seen in data files (**Supplementary Fig. 8c**). Next, we sought to answer whether chromatin changes are influencing broad categories of genes by assigning each peak to its nearest RefSeq gene TSS within 50kb. 2,345 genes were associated with more ALS peaks than control and were enriched for signaling and calcium ion binding GO terms. Conversely, 2,617 genes were associated with more control peaks than ALS and were enriched for terms such as neuron development and axon guidance (**Supplementary Fig. 8d**). Overall, ATAC-Seq identified many regulatory changes that were consistently different across ALS and control lines. In the data integration section, we analyzed how these changes correspond to changes in RNA-seq to understand differences in gene regulation between disease and control states.

### Targeted identification of potential eQTLs: WGA and RNA-Seq data integration

Our analysis of the control and ALS lines revealed genomic variants in loci other than the C9ORF72 locus that could potentially contribute to the line-specific differences in the RNA-Seq and proteomic data and influence data integration outcomes. Therefore, we evaluated whether any of the genetic coding variants outside the C9ORF72 locus were disproportionately present in C9-ALS lines compared with controls to determine whether specific genetic variants might drive expression differences between C9ORF72 and control lines that are not directly regulated by the disease mutation and could potentially confound the interpretation. While this is a very small data set and underpowered to draw significant conclusions, the goal was to establish a method and example to evaluate variants that may alter or confound the identification of signatures specifically attributable to ALS-associated HRE in C9ORF72. For example, we observed that a missense mutation in exon 17 of the poly(ADP-ribose) polymerase 1 (*PARP1*) gene (V762A) that was present in all 4 C9-ALS lines, but present in only one of the controls (**Supplementary Fig. 9**). As this was one of the genes found in the nodes of the integrated network (see below), it is possible that changes observed in the RNA-Seq data could be due to this genomic variant rather than a consequence of the HRE in C9ORF72. Further, we have no reason to believe that this variant is a haplotype that is associated with the C9ORF72 expansion. Therefore, we sought to relate the WGA to the OMICs results to better determine which genes were differentially expressed due to the HRE in C9ORF72 and which might be due to line-specific genetic variation at other loci. Although the sample size is too low for eQTL analysis, we performed this analysis to identify potential eQTL that can be reassessed using a larger sample size in future studies. The methodology we used follows standard statistical analysis for eQTL identification. We focused on exonic variants and found 7235 nonsynonymous variants that were enriched in either controls or ALS cases (**Supplementary Table 11**). Then, we compared the genes in which these variants were found to the genes that were found to be differentially expressed (FDR<0.1, which corresponds to p<0.015) in C9-ALS or control samples by RNA-Seq. We observed 801 variants (including missense, stop gain, start loss, splicing, frameshift) in genes that were differentially expressed (**Supplementary Table 11**). To examine if these subset of differentially expressed genes were significantly correlated to the presence of the variant, we performed linear regression. After voom normalization of the gene expression counts, using the limma package, a linear model was fit to each normalized gene expression-variant comparison. Adjusted R^2^ and Benjamini-Hochberg adjusted p-values were calculated for each linear fit. This linear regression analysis revealed 69 variants that could be influencing the expression of 56 genes, and confounding the identification of C9ORF72 ALS-specific gene expression differences (**Supplementary Table 12**). Seven of these genes were found in the final network analysis, but some discordance can be seen in the genotype-expression comparisons (**Supplementary Fig. 9**), which could be due to the limited number of samples for the regression analysis. To further try and determine whether genetic variants in our samples were confounding the identification of an ALS signature, we compared the variants that were enriched in either the controls or cases to known brain-specific eQTLs from the xQTL database^35^. There were 73,142 variants in our samples that overlapped with significant known brain eQTLs that represented 5292 genes; of these genes, 114 overlap genes were found to be significantly differentially expressed in the ALS versus control cases. 19 of these variants were found in all cases of one group only versus the other group, e.g. all ALS cases and no controls or no ALS case and all controls. These 19 variants are known eQTLs for 7 genes that were also found in our RNA-Seq analysis to be differentially expressed between ALS and control groups (**Supplementary Figs. 9, 10**); one of which, integrin subunit alpha V (*ITGAV*), was identified as dysregulated in each primary assay, WGA, network and as a fly modifier gene. These analyses demonstrate that the known brain eQTLs are likely to have at most a modest effect on the expression differences between C9ORF72 and control lines in our study and also provide a path for future studies with large patient cohorts.

### Comparison of RNA-Seq, proteomics and ATAC-seq experiments

We sought to characterize the similarities and differences between the genomics, RNA-Seq, proteomics, and ATAC-seq experiments. We first examined the overlap of the RNA, protein and epigenomics assays. Each differentially open region was assigned the nearest protein coding gene (up to a limit of 50kb from the TSS). The sets of genes and proteins detected by each assay all showed a modest increase in overlap compared to what would have been expected by chance. For example, approximately 7% of the proteins that differed between ALS and control samples were also differentially expressed in the RNA-Seq data (pval=1.92E-14). A higher fraction of genes that differed in RNA expression also showed changes in ATAC-seq (38%; pval=1.86E-14) and 14% of the proteins that differed between ALS and control samples were also differential ATAC-seq genes (pval=0.056). All three assays were also enriched for similar biological processes, for instance, comparison of the top GO terms from each experiment were enriched for adhesion and extracellular matrix processes, supporting biological overlap between assays (**Figure 2, Supplementary Fig. 11**).

### An “omics integrator” reveals novel C9-specific pathogenic pathways

The challenges in comparing diverse molecular assays requires a more integrated and systems-based approaches. Careful integration of the various omic data with each other and prior knowledge from the literature provide an opportunity to uncover causal relations. For example, a joint analysis of epigenomics and transcriptional data can uncover evidence of activity changes in key transcriptional regulators, which tend to be difficult to detect using mass-spectrometry. Similarly, mapping proteomic data onto networks representing protein-interactions can reveal functional relations among the differentially expressed proteins. We used a strategy implemented in Omics Integrator^36^ to explore these relationships. This approach begins by using motif analysis of open chromatin regions near differentially expressed genes to identify Omics Integrator then uses network optimization to search a vast database of protein-protein interactions to discover, *de novo*, pathways linking the experimentally determined proteomic data and the inferred transcription factors.

### Identification of transcriptional regulators

Potential transcriptional regulators were identified using *de novo* DNA motif analysis as a first step. To capture regulators mediating changes in chromatin accessibility, we searched for motifs that are enriched in differentially accessible peaks. We also searched within peaks that changed in accessibility and were near differentially expressed genes to identify transcriptional regulators that drive changes in gene expression. Peaks that were less accessible in C9-ALS samples were enriched for several TFs including Nuclear Factor I (NF1) family that controls the onset of gliogenesis in the developing spinal cord^37^ and LIM Homeobox (LHX) TFs that regulate expression of axon guidance receptors^38^ (**Figure 3a**). Conversely, peaks that were more accessible in C9-ALS samples were enriched for AP-1, RUNX2, and TEAD4. Altered AP-1 activity, which was independently predicted by IPA of the transcriptomics data, has previously been described in SOD1 mouse models^39^. Notably, we found that RNA transcripts corresponding to motifs enriched in C9-ALS peaks are upregulated in C9-ALS samples, while transcripts corresponding to motifs enriched in control peaks are downregulated in C9-ALS samples (**Figure 3b**). These results suggest that epigenetic changes could be driven by differences in expression of transcription factor transcripts.

**Figure 3:**
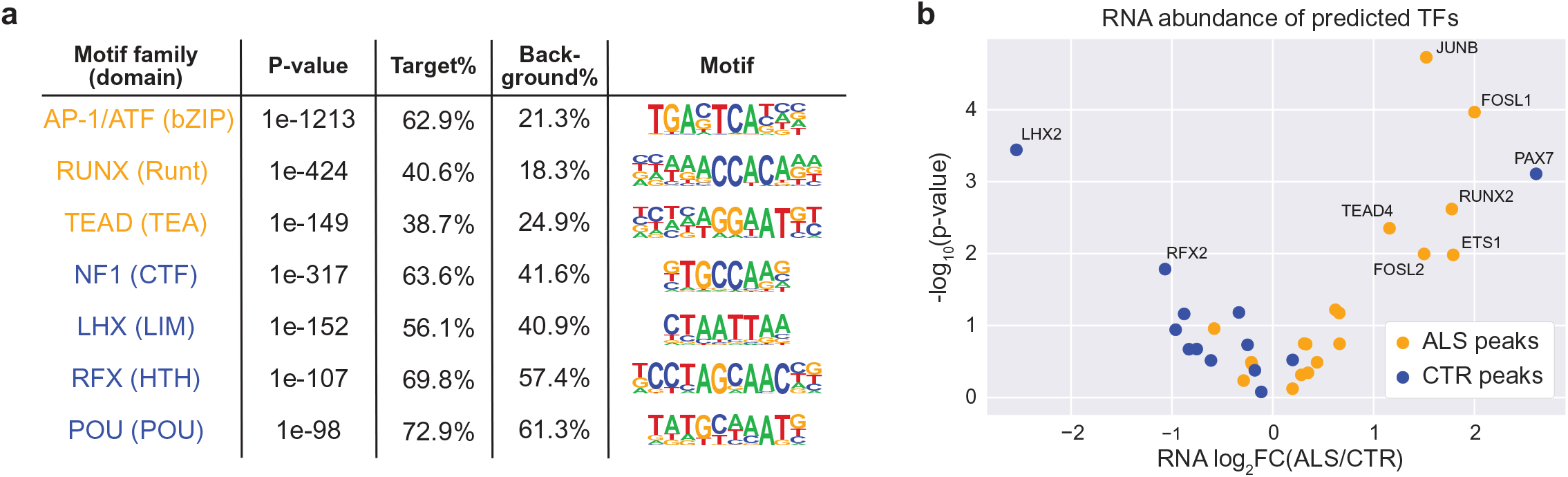
Transcription Factor predictions. (A) Transcription factor families that are predicted to be differentially active between ALS and control samples. Orange motifs are predicted to be more active in ALS and blue motifs are predicted to be more active in controls. (B) A volcano plot of RNA abundance for each predicted TF shows that TFs that are predicted to be active in ALS are also more highly expressed in ALS samples, while TFs that are predicted to be active in controls are less expressed in ALS samples.

### A network of C9ORF72-induced changes

In the next phase of the integration, we combined the transcriptional regulators inferred from RNA-Seq and ATAC-seq with the proteins detected in mass spectrometry. Our approach sought to discover, *de novo*, the cellular pathways that are differentially active between C9 and control lines. The challenge is to go beyond the limited information available in annotated pathways while still avoiding an uninterpretable network containing thousands of interactions. Our approach searches for previously reported protein-protein interactions that connect, directly or indirectly, our proteomics and transcriptional regulatory data. The method considers the strength of experimental evidence supporting each reported protein-protein interaction from the database and the strength of evidence supporting our own data.

Omics Integrator was used to search for connections among 376 predicted TFs and differentially expressed proteins. After optimization and filtering for robustness, the network retained 291 of these proteins and added 83 other proteins that were closely connected by physical interactions. The resulting 374 node network is shown in **Figure 4a**, with nodes organized by cellular compartment.

**Figure 4:**
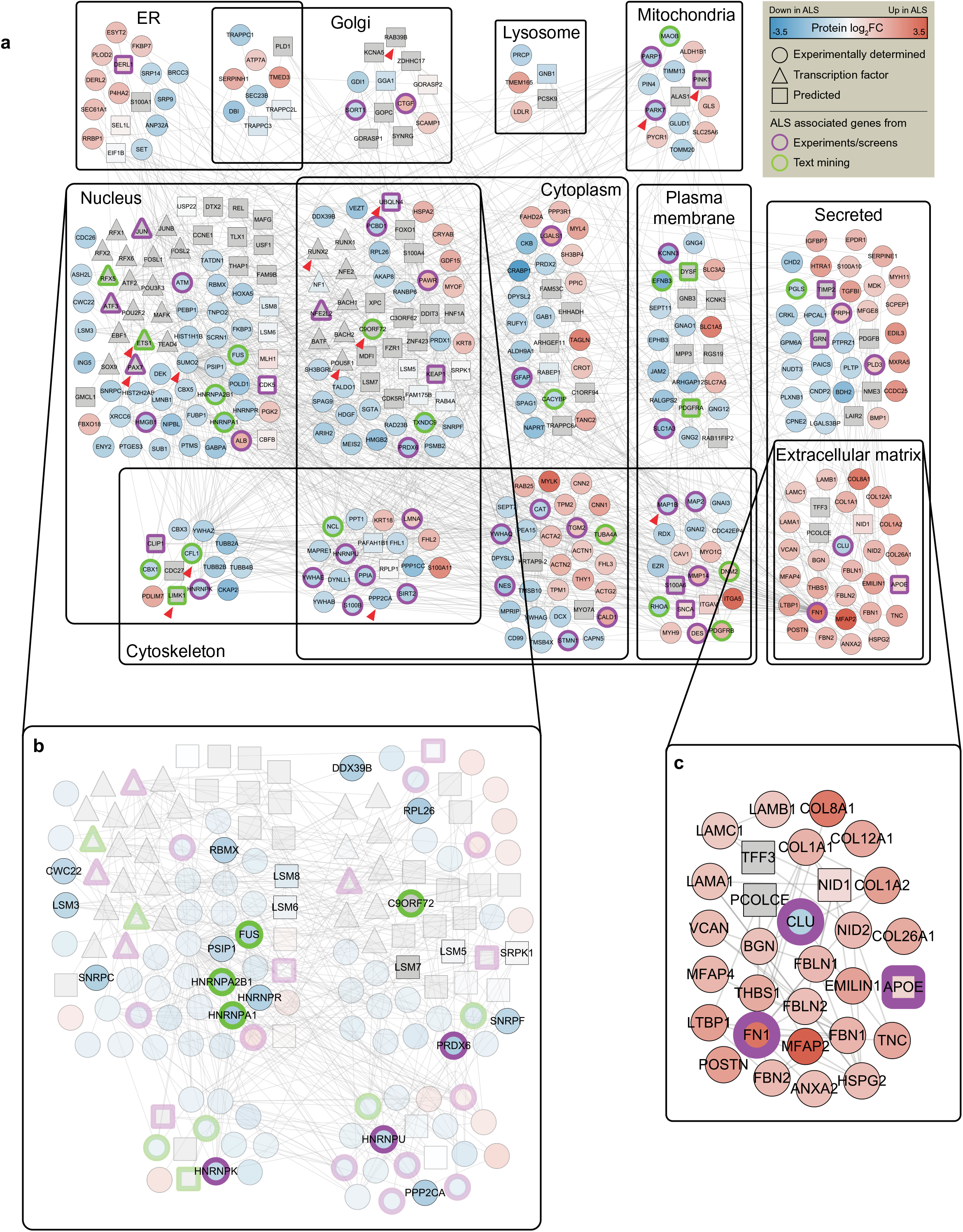
Data Integration. (A) Integrative analysis reveals a network of 374 proteins organized by subcellular location, of which 264 are experimentally determined from proteomics (circles), 27 are predicted transcription factors, and 83 are other proteins that were closely connected by physical interactions. Borders indicate ALS-associated genes from experiments or screens (purple) and text mining (green). (B) A zoomed in view of the nucleus compartment displaying genes with RNA metabolism functions. (C) A zoomed in view of the extracellular matrix compartment.

To evaluate the performance of our algorithm, we assessed the network for enrichment of genes previously associated with ALS (see **Supplementary Table 13** for ALS gene composition). We found strong enrichment for ALS-associated proteins (**Figure 4a** bolded borders; pval=4.0E-13). We also found that the 83 proteins added by Omics Integrator were also enriched for ALS associated genes (pval=2.4E-3), providing confidence that our method can predict disease-relevant proteins and pathways.

In order to understand the function of the identified network we scored it using categories from Gene Ontology. Enrichment analysis revealed significant dysregulation of ECM, in line with our transcriptomic, proteomic, and epigenomic results. Furthermore, the network was enriched for proteins belonging to cytoskeletal organization and RNA metabolism pathways (**Figure 4a,b**), both previously implicated in ALS. For instance, the nuclear-cytoskeletal compartment contains cofilin (CFL1), a known interaction partner of C9ORF72 that modulates actin dynamics in motor neurons^40^. LIMK1, a kinase that phosphorylates CFL1 also appears in the network and is known to also phosphorylate MMP14 (found in the cytoskeletal-plasma membrane compartment in **Figure 4a,c**), an endopeptidase that degrades ECM components^41^. Proteins involved in microtubule organization (PPP2CA, MAP1B, tubulin) are also represented in the cytoskeletal component of the network. PPP2CA, a major phosphatase for microtubule-associated proteins and a known binding partner of C9ORF72, has been shown to activate MAP1B which in turn tyrosinates tubulin^42^. Our network also features mitochondrial proteins that are involved in responses to oxidative stress. Mutations in PARK7 have been linked to ALS^43^, and its knockdown has been shown to increase disease severity in SOD1 mouse models^44^. Furthermore PINK1, a PARK7 mitochondrial cofactor, plays a role in axonal transport of mitochondria^45^. Lysosomal dysfunction has also been implicated in ALS^46^. Small GTPase RAB39B plays an important role in the initiation of autophagy via C9ORG72’s GDP-GTP exchange factor activity^47^. UBQLN4, linked to ALS and found in the cytoplasmic component of the network, may assist in maturation of autophagosomes^48^.

The network also reveals potentially pathological interactions between differential proteins and predicted transcriptional regulators. SUMOylation via SUMO2 is a post-translational modification process that can affect structure, localization, activity, and stability of substrates. Specifically, SUMOylation of POU5F1 (Oct4) and PAX7 enhances their stability and transactivity^49, 50^ while SUMOylation of JUN (AP1 family), ETS1, and RUNX2 reduces their stability and transactivity^51, 52^. Notably, SUMO2 protein is downregulated in ALS samples, and the activity of these transcriptional regulators following SUMOylation is concordant with their predicted activity in **Supplementary Fig. 12**. SUMOylation’s role in affecting the stability of hnRNPs and localization of actin components to the nucleus has previously been reported^53, 54^. Finally, a recent study showed that SUMOylation of stress granule (SG) proteins is required for disassembly, which is impaired by C9orf72-assicated dipeptide repeats (https://www.biorxiv.org/content/10.1101/2020.01.29.830133v1). Our analysis suggests that SUMOylation may have substantial influence on transcriptional regulation in C9-ALS motor neurons.

### Validation of integrated network in human postmortem C9 cervical spine

In order to assess the statistical and biological rigor of our data, we compared our network data to RNAseq data from an independent cohort of 12 C9 and 10 control subject postmortem cervical spine and found a significant overlap between these differentially expressed genes (3168 DEGs, at FDR<0.1) and our integrated network, especially of the ECM subnetwork (8 overlapping genes). To explore the possibility that the network optimization biased this result, we also computed an empirical p-value, as follows. Omics Integrator was run on 100 randomized inputs, generating 100 randomized networks (see methods section for details). We then computed significance (Fisher’s Exact test) of overlap between the network nodes (genes/proteins) and differentially expressed genes in the postmortem cervical spine for each randomized network and plotted in **Supplementary Fig. 13a. Supplementary Fig. 13b** shows a density plot for the number of overlapping genes from the 100 randomized networks. The mean of the distribution of p values is shifted far from our true p value. Only 1 randomized network reached a significance of enrichment greater than our true network (empirical p-value <0.01), with a large number (42) not even reaching significance at an alpha of 0.05. Additionally, using 1000 random permutations of patient condition labels for the postmortem data, we assessed the statistical significance of those DEGs (**Supplementary Fig. 13c**). Out of 1000 permutations, only 1 had a number of DEGs > 3168, indicating an empirical p-value <0.001. These 1000 sets of DEGs were then overlapped with the ECM subnet (30 genes) in the integrated network and distribution of the number of overlapping genes is shown in **Supplementary Fig. 13d**. Out of 1000 permutations, none of the DEG lists have an overlap >=8, indicating an empirical p-value <0.001. Taken together, dysregulation of the ECM subnetwork from our iPSC derived motor neuron study is supported by the human post mortem data.

### Validation of key pathways from the literature and using a fly screen

Many of the pathways identified using the Omics Integrator could also be found by searching the current ALS literature as described above. In addition, there were novel pathways that had not previously been reported as disrupted in C9-ALS including the extracellular matrix and cytoskeleton. In order to validate our “integrated omics” list generated from control and C9-ALS iMNs *in vivo*, we conducted an RNAi-based screen in a Drosophila model of G4C2-mediated neurodegeneration^55^. In this model, over-expression of 30 G4C2 repeats in the eye leads to age-dependent photoreceptor neurodegeneration, and genetic pathways identified as modifiers of fly eye degeneration have proven to be relevant to C9ORF72-associated neurodegeneration in mouse and human iPSC-derived neuron models^55, 56^. Following identification of fly homologs of human genes identified in our omics analyses (see methods), a total of 288 fly genes corresponding to 242 human genes were knocked down in the G4C2 fly model and their ability to modify (suppress or enhance) the rough eye phenotype was scored (**Figure 5b**). When available, multiple RNAi lines were tested. Of those, about 20% enhanced and 15% suppressed C9 toxicity (**Supplementary Table 14**) with a score of at least +/−1 respectively (**Supplementary Fig. 14**). The remainder showed little or no effect on eye degeneration and approximately 2% resulted in lethality. There was no particular relationship between the proteomic changes in iMNs and the phenotypic effect of knocking-down the gene in the fly (**Supplementary Table 15**). Although the precise mechanism by which these genetic manipulations affect neurodegenerative phenotypes in the fly eye at this time, the results from the fly screen confirm that a subset of genes/proteins, identified through our integrated omics approach, are capable of modifying and/or contributing to C9ORF72 G4C2-repeat-mediated toxicity.

**Figure 5:**
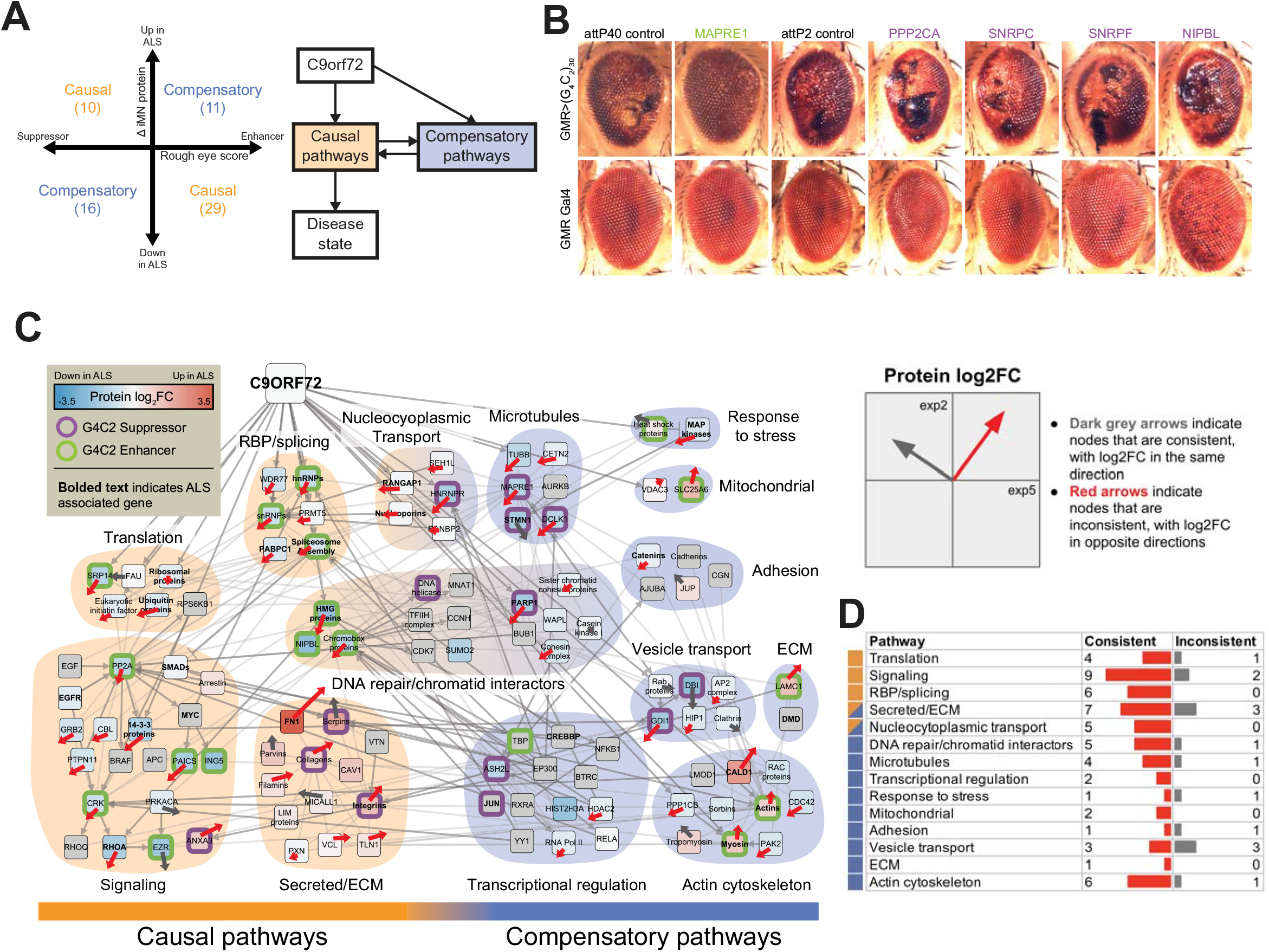
Validation in Drosophila. (A) Left: Each gene that was tested in the fly model are sorted into causal or compensatory categories using its fly phenotype and change in protein values in iMNs. Right: A schematic showing the interplay between causal and compensatory pathways that eventually result in the disease. (B) The effect of genetic manipulations on external eye morphology and depigmentation in G4C2 expressing flies. (C) Causal and compensatory genes from A were connected via intermediate genes and the resulting network was organized by cellular process. Proteins from the same families were consolidated into a single node for readability. The borders indicate whether the gene is a G4C2 suppressor (purple) or enhancer (green). Bolded names indicate ALS-associated genes. The horizontal and vertical components of the arrows indicate protein fold changes (ALS/CTR) between the original and validation experiment, respectively. Red arrows indicate proteins whose fold changes were consistent between experiments while black arrows indicate proteins that were inconsistent. (D) Numbers of consistent and inconsistent nodes in each pathway in C.

### Characterization of putatively causal and compensatory pathways

We leveraged the fly results to explore the potential causal roles of proteins that changed in the iMN data. Based on omic data alone, where specific genes, proteins and pathways are identified as up- or down-regulated, it is not possible to determine whether a difference in ALS versus control motor neurons is part of the toxic effects of the C9ORF72 expansion or whether it represents a compensatory process. However, we can begin to resolve this ambiguity using the results of the RNAi screens carried out in the fly model of the repeat expansion above. For example, in the simplest case, if a protein is upregulated in C9-ALS motor neuron cultures and knocking it down suppresses eye degeneration in the fly, the ALS-induced change(s) were likely deleterious. We refer to such C9-induced changes as “causal.” By contrast if knock-down of the same protein resulted in enhanced eye degeneration, the ALS-induced change(s) are more likely to be part of a compensatory adaptation. In total, we found 39 causal and 27 compensatory genes (**Figure 5a, and examples in Figure 5b**).

We developed an integrative approach to discover the functional interactions among these genes and their underlying roles in ALS pathology. Specifically, we built networks connecting these proteins using directed interactions gathered from two public pathway databases - KEGG and Reactome (see Methods) and grouped the resulting proteins by functional categories (**Figure 5a**).

This approach revealed several causal pathways (**Figure 5c**) that were previously known to be dysregulated by the mutated form of C9ORF72, such as RNA splicing and nuclear transport^56, 57^, the altered proteins in these pathways include ALS and associated genes such as hnRNPA1, FUS (located in the Spliceosome Assembly node), and RanGAP1 (**Supplementary Table 13**). Other pathways emerged as causal that have been less thoroughly examined in the context of C9ORF72. These pathways include signaling pathways such as EGF signaling, SMAD signaling (eg: EZR and CRK) with a hub centered around phosphatase PP2A. The approaches used here also highlighted a novel, causal set of ECM-related pathways and genes including integrins, collagens and serpins. Within these networks based on the fly data, a number of pathways are likely to represent compensatory changes. For instance, the observed increases in the cytoskeletal proteins like actin, myosin and tropomyosin, increases in heat shock proteins and decreases in RAC proteins and other proteins relating to GTP/GDP exchange are compensatory. Our approach also begins to reveal interactions between different processes. For example, the putatively causal toxic changes in the nucleocytoplasmic transport or oxidative stress are connected to potentially compensatory changes in DNA repair pathways. Finally, regulation of causal and compensatory processes can be elucidated using this approach (**Figure 5c**). As an example, while ECM/secreted proteins fall into causal pathways, cell adhesion protein changes are largely compensatory, as is dysregulation of Laminin C1, which is a component of the basal lamina and is secreted and incorporated into ECM matrices as an integral part of the structural scaffolding in tissues.

### Validation with a different motor neuron culture

We next asked whether the data integration could be validated using a replication cohort and tested our results using a different cellular model and with 6 C9orf72 patients and 7 controls, none of which were in common with the original set of samples (**Supplementary Fig. 4**). We used a modified motor neuron differentiation protocol termed direct iPSC-derived Motor Neurons 18 d diMNs^26, 58^ (**Supplementary Fig. 2**) comprised of three main stages to allow us to investigate the robustness of the original findings. In Stage 1, neural induction and hindbrain specification of iPSCs is achieved by inhibition of dual SMAD and GSK3β pathways. During Stage 2, specification of spinal motor neuron precursors is achieved by addition of Shh agonists and retinoic acid. Maturation of these precursors into neurons with more complex processes and neurites occurs during the final stage 3 with addition of neurotrophins and Notch pathway antagonists. This protocol generated an equivalent overall neuronal composition consisting of ∼75-80% of βIII-tubulin (TuJ1) positive neurons in both CTR and C9-ALS diMNs cultures (**Supplementary Fig. 2 and 3**). The percent of spinal motor neuron positive cells marked by SMI32 (NEFH) and NKX6.1 positive cells were also not statistically different between CTR and C9-ALS cultures – SMI32 (CTR = 59.7% and C9-ALS = 43.5%) and NKX6.1 (CTR = 39.3% and C9-ALS = 27.1%). However, a significantly different percent of ISLET1 positive cells was observed in control (41.1%) versus C9-ALS (26.9%) cultures (pval=0.0009) in day 18 of diMNs at the time point of cell harvest (**Supplementary Fig. 2 and 3**).

RNA-seq and proteomics were then performed on each of the samples. The differential signals for the original and validation experiments were correlated for RNA (pearson=0.25; pval=8.9E-224) and proteomics (pearson=0.24; pval=1.7E-33). We identified 91 differential transcripts in RNA-seq which were enriched for actin cytoskeleton terms, similar to the enrichments from the original analysis. Of these 91 transcripts, 6 were in common with the original set of differential genes (pval=0.0015). Proteomics also identified 250 differential proteins that were enriched for ECM, adhesion and axon related terms (adhesion and collagen are not currently highlighted in the draft, but we should), though RNA binding and splicing terms were not found to be enriched. Of these 250 proteins, 68 were in common with the original set of differential genes (pval=0.05).

Next, we explored how well the validation data confirmed our original integrated network (**Figure 4**) by comparing the signed proteomics log fold change for each node in the network with proteomics from the validation data. Proteins were significantly more likely to be consistent between experiments, with 133 proteins changing in the same direction versus 40 proteins changing in opposite directions (pval=1.5E-12). More secreted/ECM proteins were consistent than not (11 versus 1) and more cytoskeletal proteins were consistent than not (40 versus 12) (**Figure 5d**, **Supplementary Fig. 15**). The differential protein expression for nodes in the network were significantly correlated (pearson=0.49; pval=5.5E-12) indicating good concordance between the original and validation data.

## Discussion

iPSC models offer a way to map the initiation and execution of pathology in specific diseases of the central nervous system (CNS). This is clearly required given the lack of effective drugs for brain disorders despite years of investment from both industry and academia. Many groups have now been able to generate iPSCs from patients with neurological disease-causing mutations and have shown specific phenotypes in the dish^59, 60^ and we recently reviewed the many studies using iPSCs to model ALS^58^. More recent studies show a stress-induced phenotype in C9 iPSC-derived motor neurons^17^ and an overall cell death and reduced fiber outgrowth phenotype in a range of ALS cases not including C9ORF72^19^. In another report, increased activity in motor neurons from ALS patients in the dish led to a drug trial with retigabine^61^. Interestingly, all of these studies were focused on discovering physical *in vitro* phenotypes such as cell death or reduced fiber outgrowth, which may or may not be relevant to drug intervention in patients. One of the key difficulties in these studies has been an incomplete picture of the earliest and most significant changes that occur during pathogenesis.

Based on the premise that dysfunction of molecular pathways in specific cell populations in the brain leads to neurodegeneration, we have established a comprehensive, quantitative molecular phenotyping approach using a human iPSC technology platform to study molecular signatures of CNS cell types focusing on iPSCs from patients with C9orf72, given its prevalence as a genetic cause of ALS and its dominant phenotype^5^. We have used genomics, transcriptomics, epigenomics, and high-content, quantitative proteomics to characterize human motor neuron cultures from C9orf72 ALS patients, under strict quality control including the use of *parallel and identical* cultures for each assay, metadata standards, and analytical pipelines. The use of the same cultures was critical, given that cell type heterogeneity arising from iPSC differentiations and batch effects can be a confound in molecular analyses of long-term differentiations^62^. Given that the C9 mutation is a variable sized intron expansion of G4C4 and that all unaffected people normally have variable numbers of repeats as well, the generation of isogenic lines that replace the expansion with a “normalized” repeat has been challenging and is not practical to ultimately evaluate large numbers of sporadic disease patients, therefore we used lines derived from different patients.

A computational pipeline was used to integrate the diverse molecular data sets and identify the most significant regulated pathways in patient cells. This “OMICS Integrator” software^36^ uses network approaches to integrate diverse data types into coherent biological pathways that can avoid some of the pitfalls associated with analyzing single data types and uncover novel pathways that are not annotated in existing databases. This approach is validated by the strong statistical enrichment and the comprehensive number of hits it recovered that are consistent with published literature for C9orf72 ALS. The significant overlap between our integrated network, especially of the ECM subnetwork and data from C9orf72 ALS postmortem cervical spine tissues, further validates the significance of the networks identified. At the same time, the approach revealed functional links among the disparate data, including identifying many transcriptional regulators.

A challenge in using multi-omic data sets is understanding how the direction of a change impacts disease pathogenesis. This is perhaps one of the greatest difficulties – e.g. understanding if the observed changes are conducive to the course of the disease or a cellular attempt at a homeostatic response to physiological insults. Using Drosophila genetics guided by the outcome of the integrated networks, it has been possible to not only validate the specific genes and proteins involved, but also to discern probable effect and whether altered expression or activity would be predicted to promote disease pathogenesis or serve as a compensatory response. The results of these studies provide a unique data source and methods that can be utilized in the study of ALS and other neurodegenerative diseases.

Our analysis reveals a complex system of interweaving relationships among causal and compensatory pathways. In some cases, such as the ECM, causal and compensatory roles were found to exist even within the same pathways. Though the literature on the ECM’s role in neuronal function and disease progression is limited, several studies have described neuroprotective properties of ECM^63^ and a proteomic study of ALS subject CSF revealed the ECM as an enriched biological process^64^. Our analysis suggests that, while ECM components are broadly upregulated in ALS, individual components of the ECM may have very different downstream consequences. For example, knocking down some proteins like LAMC1 and DMD enhances toxicity in fly eyes while knocking down other ECM components like Serpins, Collagens and Integrins suppress toxicity. One mechanism through which extracellular signals within the ECM may be internalized is through integrin signaling. Integrin activation mediates molecular coupling of CAS and Crk, and the resulting complex has been shown to regulate the actin cytoskeleton^65^. Interestingly, integrins and CRK were both found to be pathogenic, while actin cytoskeletal components were compensatory, which suggests ECM pathogenicity is transmitted via some non-cytoskeletal pathway.

It is also important to recognize that the classification of changes as “causal” or “compensatory” is far from definitive. Not all results from the fly necessarily translate to human cells and tissue. Furthermore, our simple binary classification does not capture complicated situations in which there may be non-linear effects of gene expression on phenotypes. However, these first attempts at relating many different aspects of cell functioning are the starting blocks for further studies and enable for the first time the development of a holistic view of cell functioning in the face of a pathogenic repeat that causes ALS.

Studies of human derived samples must account the variation that can result from individual variation as well as differences in cell state that may not be disease related. In our study, we separately examined cells from two different cohorts that were differentiated using different protocols. While the details of the differentially expressed genes and proteins varied across the cohorts, our integrative approach was able to highlight many systems-level similarities. It is worth noting that large differences between cohorts is just as much an issue in studies of post-mortem tissue as it is in studies of iPSC-derived material. For example, a recent proteomic analysis of Alzheimer’s post-mortem brain samples found that of 173 proteins initially detected as differentially expressed in brain, only 58 showed consistent changes in a second cohort, and only 34 of these showed consistent changes in RNA levels^66^. Focusing on network-level changes may help uncover commonalities that transcend differences among samples.

This integrative approach is well suited for the task of hypothesis generation. For instance, our results suggest that DNA repair pathways are a compensatory response to either nucleocytoplasmic transport deficits or oxidative stress. In addition to providing insight into how these pathways interact, our analysis also identifies proteins that are attractive targets such as MAPRE1. While we acknowledge there are some limitations of integrating data across human *in vitro* and fly *in vivo* models, this approach provides a much needed basis for establishing causality and generating testable hypotheses.

An additional benefit in having transcriptomic and proteomic data together with WGS is the ability to integrate these data sets and identify whether a given DNA sequence change causes altered expression of the gene or altered levels of the protein. Using the data set here, we have integrated WGS with RNASeq data to establish a pipeline and begin to evaluate eQTLs that may be meaningful to disease as a causal modifier versus altering gene expression as a consequence of ALS. Future studies will expand this analysis across each of assays and extend to larger data sets from additional ALS subjects.

Based on hypotheses generated through the generation of integrated networks and potential causality suggested by the fly data, next steps will include testing whether modulation of these pathways in the iPSC neurons can impact key pathogenic features of C9orf72 such as the nuclear pore deficit and formation of dipeptide repeats. Using imaging platforms such as longitudinal robotic imaging, there is now the potential to use reporters to query specific networks or processes (e.g. ECM) in future studies^67^. The goal of this study was to establish a platform to evaluate multiple subjects and incorporate human variation by gathering and integrating a wide range of critical information using highly systematic approaches. This study was underpowered with regard to numbers of patients and made no connection to the complex clinical course of the disease. Currently we are producing 1000 iPSC lines from patients with all types of ALS (including C9orf72 mutation carriers) and performing a similar analysis under the auspices of Answer ALS (https://www.answerals.org/). In addition, the clinical history of each patient will be combined with the Omics Integrator and other computational approaches to give more resolution on how molecular changes may impact the clinical course of the disease. However, the core techniques and integrated approach of the current report along with the first set of data suggesting a molecular signature for C9 ALS provide a strong framework for this new “big data” approach to learning more about the causes and treatments of diseases such as ALS.

## Acknowledgements

We thank the ALS patients and their families for their essential contributions to this research. We also thank Dr. Shana Svendsen for editorial assistance. We also acknowledge and thank the Target ALS Postmortem Tissue Core for their contribution of samples and data. **Funding:** Primary support for this work was from NIH U54 NS091046 NeuroLINCS center (S.F., E.F., J.D.R., C.N.S., J.V.E., L.M.T.). Additional support was provided by NIH NS089076 (L.M.T, E.F), NS085207, NS094239, Fidelity Bioscience (J.D.R), The Robert Packard Center for ALS Research at Johns Hopkins, Answer ALS Project. The WGA was funded by the ALS Association and conducted at the New York Genome Center. The sequencing activities at NYGC were supported by the ALS Association (ALSA) and The Tow Foundation. This work was made possible, in part, through access to the Genomic High Throughput Facility Shared Resource of the Cancer Center Support Grant (CA-62203) at the University of California, Irvine.

## Author Contributions

<u>Designed the experiments</u>: DS, CNS, RGL, LMT, VD, JVE, PM, MA, JL, EF, JK, SF, ANC, JGD, TEL, JDR

<u>Cell Lines</u>:

Generated iPSC lines in study: LO, EG, LP, DS

iPSC culture and neuronal differentiation: BM, HT, MGB, BS

<u>Carried out experiments</u>: BM, HT, MGB, BS: differentiation; RGL, J.S.: RNA-Seq; VD, JVE PM: proteomics, MA, BTW: ATAC-Seq;

<u>Analyzed the data</u>: RGL, JW, MC, LMT; VD, JVE; PM, MA, BTW, JL, RE-C, AL, KS, NL P-M, DR, CNS, JAK, LL, SW, SF

<u>Wrote the manuscript</u>: DS, CNS, RGL, JW, LMT; CD, JVE, JL, EF, ANC, JDR, JAK, SF

<u>Project Leadership and Management</u>: SF, EF, JDR, DS, CNS, JVE, LMT, LO, TT

## Supplemental Author List

**NYGC ALS Consortium**

Hemali Phatnani, PhD

Center for Genomics of Neurodegenerative Disease (CGND), New York Genome Center, New York, NY

Justin Kwan, MD

Department of Neurology, Lewis Katz School of Medicine, Temple University, Philadelphia, PA

Dhruv Sareen, PhD

Cedars-Sinai Department of Biomedical Sciences, Board of Governors Regenerative Medicine Institute and Brain Program, Cedars-Sinai Medical Center, and Department of Medicine, University of California, Los Angeles, CA

James R. Broach, PhD

Department of Biochemistry and Molecular Biology, Penn State Institute for Personalized Medicine, The Pennsylvania State University, Hershey, PA

Zachary Simmons, MD

Department of Neurology, The Pennsylvania State University, Hershey, PA

Ximena Arcila-Londono, MD

Department of Neurology, Henry Ford Health System, Detroit, MI

Edward B. Lee, MD, PhD

Department of Pathology and Laboratory Medicine, Perelman School of Medicine, University of Pennsylvania, Philadelphia, PA

Vivianna M. Van Deerlin, MD, PhD

Department of Pathology and Laboratory Medicine, Perelman School of Medicine, University of Pennsylvania, Philadelphia, PA

Neil A. Shneider, MD, PhD

Department of Neurology, Center for Motor Neuron Biology and Disease, Institute for Genomic Medicine, Columbia University, New York, NY

Ernest Fraenkel, PhD

Department of Biological Engineering, Massachusetts Institute of Technology, Cambridge, MA

Lyle W. Ostrow, MD, PhD

Department of Neurology, Johns Hopkins School of Medicine, Baltimore, MD

Frank Baas, MD, PhD

Department of Neurogenetics, Academic Medical Centre, Amsterdam and Leiden University Medical Center, Leiden, The Netherlands

Noah Zaitlen, PhD

Department of Medicine, Lung Biology Center, University of California, San Francisco, CA

James D. Berry, MD, MPH

ALS Multidisciplinary Clinic, Neuromuscular Division, Department of Neurology, Harvard Medical School, and Neurological Clinical Research Institute, Massachusetts General Hospital, Boston, MA

Andrea Malaspina, MD, PhD

Centre for Neuroscience and Trauma, Blizard Institute, Barts and The London School of Medicine and Dentistry, Queen Mary University of London, London, and Department of Neurology, Basildon University Hospital, Basildon, United Kingdom

Pietro Fratta, MD, PhD

Institute of Neurology, National Hospital for Neurology and Neurosurgery, University College London, London, United Kingdom

Gregory A. Cox, PhD

The Jackson Laboratory, Bar Harbor, ME

Leslie M. Thompson, PhD

Department of Psychiatry & Human Behavior, Department of Neurobiology and Behavior

University California, Irvine, CA

Steve Finkbeiner, MD, PhD

Taube/Koret Center for Neurodegenerative Disease Research, Roddenberry Center for Stem Cell Biology and Medicine, Gladstone Institute

Efthimios Dardiotis, MD, PhD

Department of Neurology & Sensory Organs, University of Thessaly, Thessaly, Greece

Timothy M. Miller, MD, PhD

Department of Neurology, Washington University in St. Louis, St. Louis, MO

Siddharthan Chandran, PhD

Centre for Clinical Brain Sciences, Anne Rowling Regenerative Neurology Clinic, Euan MacDonald Centre for Motor Neurone Disease Research, University of Edinburgh, Edinburgh, United Kingdom

Suvankar Pal, MD

Centre for Clinical Brain Sciences, Anne Rowling Regenerative Neurology Clinic, Euan MacDonald Centre for Motor Neurone Disease Research, University of Edinburgh, Edinburgh, United Kingdom

Eran Hornstein, MD, PhD

Department of Molecular Genetics, Weizmann Institute of Science, Rehovot, Israel

Daniel J. MacGowan, MD

Department of Neurology, Icahn School of Medicine at Mount Sinai, New York, NY

Terry Heiman-Patterson, MD

Center for Neurodegenerative Disorders, Department of Neurology, the Lewis Katz School of Medicine, Temple University, Philadelphia, PA

Molly G. Hammell, PhD

Cold Spring Harbor Laboratory, Cold Spring Harbor, NY

Nikolaos. A. Patsopoulos, MD, PhD

Computer Science and Systems Biology Program, Ann Romney Center for Neurological Diseases, Department of Neurology and Division of Genetics in Department of Medicine, Brigham and Women’s Hospital, Boston, MA, Harvard Medical School, Boston, MA, Program in Medical and Population Genetics, Broad Institute, Cambridge, MA

Oleg Butovsky, PhD

Ann Romney Center for Neurologic Diseases, Brigham and Women’s Hospital, Harvard Medical School, Boston, MA

Joshua Dubnau, PhD

Department of Anesthesiology, Stony Brook University, Stony Brook, NY

Avindra Nath, MD

Section of Infections of the Nervous System, National Institute of Neurological Disorders and Stroke, NIH, Bethesda, MD

Robert Bowser, PhD

Department of Neurology, Barrow Neurological Institute, St. Joseph’s Hospital and Medical Center, Department of Neurobiology, Barrow Neurological Institute, St. Joseph’s Hospital and Medical Center, Phoenix, AZ

Matt Harms, MD

Department of Neurology, Division of Neuromuscular Medicine, Columbia University, New York, NY

Eleonora Aronica, MD, PhD

Department of Neuropathology, Academic Medical Center, University of Amsterdam, Amsterdam, The Netherlands

Mary Poss, DVM, PhD

Department of Biology and Veterinary and Biomedical Sciences, The Pennsylvania State University, University Park, PA

Jennifer Phillips-Cremins, PhD

New York Stem Cell Foundation, Department of Bioengineering, School of Engineering and Applied Sciences, University of Pennsylvania, Philadelphia, PA

John Crary, MD, PhD

Department of Pathology, Fishberg Department of Neuroscience, Friedman Brain Institute, Ronald M. Loeb Center for Alzheimer’s Disease, Icahn School of Medicine at Mount Sinai, New York, NY

Nazem Atassi, MD

Department of Neurology, Harvard Medical School, Neurological Clinical Research Institute, Massachusetts General Hospital, Boston, MA

Dale J. Lange, MD

Department of Neurology, Hospital for Special Surgery and Weill Cornell Medical Center, New York, NY

Darius J. Adams, MD

Medical Genetics, Atlantic Health System, Morristown Medical Center, Morristown, NJ, Overlook Medical Center, Summit, NJ

Leonidas Stefanis, MD, PhD

Center of Clinical Research, Experimental Surgery and Translational Research, Biomedical Research Foundation of the Academy of Athens (BRFAA), 4 Soranou Efesiou Street, 11527, Athens, Greece;

1st Department of Neurology, Eginition Hospital, Medical School, National and Kapodistrian University of Athens, Athens, Greece

Marc Gotkine, MD

Neuromuscular/EMG service and ALS/Motor Neuron Disease Clinic, Hebrew University-Hadassah Medical Center, Jerusalem, Israel

Robert H. Baloh, MD. PhD

Board of Governors Regenerative Medicine Institute, Los Angeles, CA;

Department of Neurology, Cedars-Sinai Medical Center, Los Angeles, CA

Suma Babu, MBBS, MPH

Neurological Clinical Research Institute, Massachusetts General Hospital, Boston, MA

Towfique Raj, PhD

Departments of Neuroscience, and Genetics and Genomic Sciences, Ronald M. Loeb Center for Alzheimer’s disease, Icahn School of Medicine at Mount Sinai, New York, NY

Sabrina Paganoni, MD, PhD

Harvard Medical School, Department of Physical Medicine & Rehabilitation, Spaulding Rehabilitation Hospital, Boston, MA

Ophir Shalem, PhD

Center for Cellular and Molecular Therapeutics, Children’s Hospital of Philadelphia, Philadelphia, PA;

Department of Genetics, Perelman School of Medicine, University of Pennsylvania, Philadelphia, PA

Colin Smith, MD

Centre for Clinical Brain Sciences, University of Edinburgh, Edinburgh, UK;

Euan MacDonald Centre for Motor Neurone Disease Research, University of Edinburgh, Edinburgh, UK

Bin Zhang, PhD

Department of Genetics and Genomic Sciences, Icahn Institute of Data Science and Genomic Technology, Icahn School of Medicine at Mount Sinai, New York, NY

University of Maryland Brain and Tissue Bank and NIH NeuroBioBank

Brent Harris, MD, PhD

Department of Neuropathology, Georgetown Brain Bank, Georgetown Lombardi Comprehensive Cancer Center, Georgetown University Medical Center, Washington DC

Iris Broce, PhD

Neuroradiology Section, Department of Radiology and Biomedical Imaging, University of California, San Francisco, San Francisco, CA

Vivian Drory, MD

Neuromuscular Diseases Unit, Department of Neurology, Tel Aviv Sourasky Medical Center, Sackler Faculty of Medicine, Tel-Aviv University, Tel-Aviv, Israel

John Ravits, MD

Department of Neuroscience, University of California San Diego, La Jolla, CA

Corey McMillan, PhD

Department of Neurology, University of Pennsylvania Perelman School of Medicine, Philadelphia, PA

Vilas Menon, PhD

Department of Neurology, Columbia University Medical Center, New York, NY

Lani Wu, PhD

Department of Pharmaceutical Chemistry, University of California San Francisco, San Francisco, CA

Steven Altschuler, PhD

Department of Pharmaceutical Chemistry, University of California San Francisco, San Francisco, CA

## Competing financial interests

The other authors declare no competing interests.

## STAR Methods

### Generation and characterization of iPSC lines

The initial 3 control lines (termed 25iCTR, 83iCTR, 00iCTR) and 4 iPSC lines (termed 29iALS, 52iALS, 30iALS, 28iALS) were generated using episomal plasmids and characterized as previously described^15^. Fibroblasts from C9ORF72 ALS patients (28iALS-n2, 29iALS-n1, 30-iALS-n1, and 52iALS-n6) were derived at Washington University of St. Louis. Healthy control fibroblasts (00iCTR: GM05400; 83iCTR: GM02183) were obtained from the Coriell Institute for Medical Research. The Coriell Cell Repository maintains the consent and privacy of the donor of fibroblast samples. All the cell lines and protocols in the present study were carried out in accordance with the guidelines approved by institutional review boards at the Cedars-Sinai Medical Center and Washington University at St. Louis. Studies were performed under the auspices of the Cedars-Sinai Medical Center Institutional Review Board (IRB) approved protocol Pro00028662 and Pro00028515. The reprogramming and characterization of iPSC cell lines and differentiation protocols in the present study were carried out in accordance with the guidelines approved by Stem Cell Research Oversight committee (SCRO) and IRB, under the auspices of IRB-SCRO Protocols Pro00032834 (iPSC Core Repository and Stem Cell Program), Pro00024839 (Using iPS cells to develop novel tools for the treatment of SMA) and Pro00027006 (Cell and Tissue Analysis for Neurologic Diseases; Robert Baloh). Appropriate informed consents were obtained from all the donors. Additional iPSC lines for the replicate cohort were generated from 7 healthy controls (CS-002, LBC360179, W15-C201, W15-C206, NEUMN061ATZ, NEUVW301WP3, NEUPW469XH7) and 6 ALS patients (NEUEM720BUU, NEUVX902YNL, NEUUL256UC9, NEUPK546ZLD, NEUFV237VCZ, NEUDT709YHN) were reprogrammed from PBMCs. All lines were reprogrammed by nucleofecting parent cells with nonintegrating oriP/EBNA1 plasmids, which allowed for episomal expression of reprogramming factors similar to previously published protocols^60, 68^. All of the iPSC lines described in this study are available from the iPSC Core at the Cedars-Sinai Biomanufacturing Center iPSC repository. To protect donor privacy and confidentiality, all samples were coded and de-identified in this study. Extensive quality control processes were implemented, including testing iPSC precursors and final neuronal samples (motor neuron cultures) for purity and their identity by short-tandem repeat (STR) analysis performed by a third-party company before the samples were distributed. G-band karyotyping was performed to ensure that iPSCs maintained normal karyotypes. The parental tissue (fibroblasts and blood), reprogrammed iPSCs and the differentiated iMNs and diMNs prior to performing assays were submitted to IDEXX BioResearch for DNA fingerprinting and STR analysis to confirm donor identity. Each of the iPSC lines used in this study had unique genetic profiles and the profiles of the samples and their source tissues were identical. Additionally, the test confirmed the samples to be of human origin and detected no mammalian interspecies contamination. The Cedars-Sinai iPSC Core Facility created a working cell bank of iPSC-derived motor neuron precursor spheres (iMPS) and terminal diMNs for C9-ALS and control subjects.

### Whole Genome Sequencing and Analysis

DNA was extracted from iPSC lines using the QIAamp DNA Blood mini Kit (Qiagen; 51104) as per the manufacturer’s instructions. A minimum of 1μg of unamplified, high molecular weight, RNase treated DNA with absorbance values of OD260/280 1.7-2.0 and OD260/230 >2.0, was sent to The New York Genome Center for sequencing on the Illumina X10. Sequence data was processed on NYGC automated pipeline. Paired-end 150 bp reads were aligned to the GRCh37 human reference using the Burrows-Wheeler Aligner (BWA-MEMv0.7.8) and processed using the GATK best-practices workflow that includes marking of duplicate reads by the use of Picard tools(v1.83, http://picard.sourceforge.net), local realignment around indels, and base quality score recalibration (BQSR) via Genome Analysis Toolkit (GATK v3.4.0) (McKenna, Hanna et al. 2010; DePristo, Banks et al. 2011)(New York Genome Center).

The variant calls from NYGC were assessed by examining the actual reads for alignment issues and spot-checking the BAM files for specific variants in IGV and assessed they were of good quality. The VCFs were converted into GVCFs and performed custom annotation and intersected a subset of the omics data (RNA-Seq, ATAC Cluster) with the WGS data.

The annotation pipeline was customized to incorporate elements from ANNOVAR^69^ and KGGseq^70^ and used to generate a report, including genotypes, for each sample. These reports are available upon request. The following annotation was used: For genes and exonic variants that have clinical significance, we incorporated the Clinical Genomic Database (CGD)^71^, the Online Mendelian Inheritance in Man (OMIM)^72^, ClinVar^73^ and genes listed in the American College of Medical Genetics and Genomics (ACMG)^74^ database as well. Intervar, which is based upon the ACMG and AMP standards and guidelines for interpretation of variants, was also incorporated. This tool uses 18 criteria to assess the clinical significance of variants and classify them based on a five-tiered system^28^. To flag ALS genes, we incorporated ALS gene lists and variants from ALSoD^75^ (http://alsod.iop.kcl.ac.uk/), a highly curated list from Dr. John Landers and ALS associations from the DisGeNet database^76^. We also incorporated functional prediction by using *in silico* prediction from nine programs, including the databases, such as SIFT^77^, PolyPhen2^78^, and Mutation Taster^79^ and as in Li et al., 2013^80^ for each variant. As well, additional databases were included that assess the variant tolerance of each gene using the RVIS^81^, the Gene Damage Index (GDI)^82^ and LoFTool^83^. Gene expression: For variants in genes that are highly expressed in the brain, we provided these data from the Human Protein Atlas^84^ (http://www.proteinatlas.org) and expression data for the cortex and spinal cord from the GTEx portal^85, 86^ (https://gtexportal.org/home/). Frequency information came from three databases on all known variants from ExAC^87^, the NHLBI Exome Sequencing Project (ESP)^88^, and the 1000 Genomes Project^27^.

A separate annotation pipeline was developed for variants that are in intergenic and regulatory regions. We report the variant in relation to the closest gene, and are either intronic, upstream, downstream (up to 4 KBs from the start and stop of a gene) or in 5’ or 3’ UTRs. The annotation used came from RegulomeDB, which annotates variants with known or predicted regulatory elements such as transcription factor binding sites (TFBS), eQTLs, validated functional SNPs and DNase sensitivity^89^. The source data comes from ENCODE^90, 91^ and GEO^92^. We also included other regulatory databases, such as Target Scan, which is an algorithm that uses 14 features to predict and identify microRNA target sites within mRNAs^93^ and miRBase^94–96^.

### Differentiation of iPSCs into Motor Neurons

#### Initial Cohort

Control and ALS iPSCs were differentiated into motor neurons (iMNs) based on a combination of previous models established for rapid neural differentiation (**Figure 1a, Supplementary Fig. 1**)^58^. Briefly, iPSCs were grown to near confluence devoid of spontaneous differentiation under normal maintenance conditions prior to the start of differentiation. Neuroectoderm specification of iPSCs was induced by removal of mTeSR1 media and addition of defined neural differentiation media (NDM) +LS composed of IMDM supplemented with B27 + vitamin A (2%), N2 (1%), Non-Essential Amino Acids (NEAA, 1%) and penicillin-streptomycin-amphotericin (PSA, 1%) along with LDN193189 and SB431542 [LS] - as a combination of small molecule inhibitors of SMAD pathway, BMP type 1 receptors (ALK2/3) TGF-beta superfamily type 1 activin receptor-like kinase (ALK) receptors (ALK4/5/7)] (**Supplementary Fig. 1a**). Colonies were dissociated into single cells with Accutase and uniform aggregates were formed in sterilized V-bottom 384-well PCR plates with 20,000 cells/well. Uniform neural aggregates were formed by seeding in NDM+LS in presence of Matrigel and centrifuging for 5 minutes at 200*g*. The aggregates were maintained in this media for 5 days. The culture medium was replenished every 2 days. On day 5, the aggregates were gently isolated from the plates using Accutase, and the uniform sized neural aggregates were then plated on laminin-coated 6-well plates. After 7 days (day 12), media were changed to a motor neuron specification medium (MNSM) generating caudo-ventralized MN precursors by addition of all-trans retinoic acid (ATRA) and the sonic hedgehog agonist, purmorphamine (PMN), brain-derived neurotrophic factor (BDNF), glial cell line-derived neurotrophic factor (GDNF), ascorbic acid (AA) and dibutyryl cyclic adenosine monophosphate (db-cAMP). Over the next 4 to 8 days, neural rosettes formed and were lifted at day 16 to 20 and subsequently cultured in suspension low-attachment flasks for a further 8 days. Selected rosettes were switched to a motor neuron precursor expansion media (MNPEM) containing ATRA, PMN, and the mitogens epidermal growth factor (EGF) and fibroblast growth factor (FGF2). After an initial 8 days in the expansion medium, theinduced motor neuron precursor spheres (iMPS) were further expanded by weekly chopping for 5 weeks (passages) and cryopreserved prior to initiation of terminal differentiation stage. These iMPS were cryopreserved into aliquots for later generation of iMPS-derived motor neurons (iMNs) for OMIC analysis or to send to imaging centers for cell death assays and live-cell imaging. In order to induce terminal motor neuron differentiation, the iMPS were fully dissociated with Accutase and seeded on laminin-coated 6-well plates, and matured in Stage 1 motor neuron maturation medium (MNMM Stage 1) consisting of NDM supplemented with ATRA (0.1 μM), PMN (1 μM), db-cAMP (1 μM), ascorbic acid (AA; 200 ng/ml), Notch signaling γ-Secretase Inhibitor, DAPT (2.5 μM), BDNF; 10 ng/ml and GDNF; 10 ng/ml for 7 days (**Supplementary Fig. 1b**). Then cultures were switched to maturation medium stage 2 (MNMM Stage 2) containing Neurobasal, 1% NEAA, 1% N2, 0.5% GlutaMax, db-cAMP (1 μM), ascorbic acid (AA; 200 ng/ml), BDNF; 10 ng/ml and GDNF; 10 ng/ml for another 14 days. Mature iMN cultures were harvested and screened at 21-days post plating. These conditions allowed for motor neuron differentiation under serum-free conditions. All differentiating cultures were maintained in humidified incubators at 37°C (5% CO2 in air).

#### Replication Cohort

A second differentiation method called “the direct induced motor neuron (diMN) protocol” which comprises three stages (**Supplementary Figure 2a**) was used for the replication cohort studies. At the outset of Stage 1, plates from each iPSC line were washed with 1mL DPBS (Corning 21-031-CV)/well and then incubated in 1mL Accutase (EMD Millipore SCR005)/well for 5 minutes at 37°C. After incubation, 1mL DPBS/well was added, cells were quickly collected into multiple 15mL conical tubes and centrifuged at 161 g for 2 minutes. Each pellet was re-suspended in mTeSR and cell viability and concentration were determined by automated cell counting (Nexcelom Auto 2000), and multiple matrigel-coated 6-well plates were seeded at a density of 5×e^5^ cells/well in 2mL mTeSR media/well. Twenty-four hours following plate-down, mTeSR media was exchanged for Stage 1 media (**Table S1** for media composition). Stage 1 media was exchanged daily until Day 6. On day 6, stage 2 of the differentiation process began when, for each cell line, all wells were washed with 1mL DPBS/well and incubated in 1mL Accutase per well for 5 minutes at 37°C. After incubation, 1mL DPBS/well was added, cells were quickly collected into multiple 15mL conical tubes and centrifuged at 161 g for 2 minutes. Cells were re-suspended in Stage 2 Platedown Media (ST2PD, **Table S2** for composition), viability and cell counts determined and multiple Matrigel-coated 6-well plates were seeded at a density of 7.5×e^5^ cells/well in 2mL ST2PD/well. Twenty-four hours following platedown, St2PD was exchanged for Stage 2 media (**Table S3** for composition). Stage 2 media were exchanged every other day until day 12. On day 12 began Stage 3 of differentiation. For each cell line, Stage 2 media were completely aspirated from all wells and replaced with 2mL Stage 3 media/well. Stage 3 media (**Table S4** for composition) were exchanged every other day until Day 18 of differentiation. During feedings, approximately 75% of old media were aspirated and 2mL Stage 3/well were added dropwise in a circular manner in order to minimize disruption of the cell monolayer. On Day 18 of differentiation, cell lines were collected and pelleted. Prior to collection and pelleting, one 6-well plate or a 96-well plate seeded in parallel and carried through the entire protocol was selected from each line for immunocytochemistry-based quantification of select motor neuron and pan neuron markers including SMI32 (NEFH), Islet1, Nkx6.1, and TUJ1 (TUBB3) (Molecular Devices ImageExpress Micro) (**Supplementary Fig. 2b, 3**). Multiple regions (9–16) of interest were captured per well for four wells at a magnification of 10×. After imaging, the plates were collected with their respective lines.

Multiple wells/cell line were set aside for short tandem repeat (STR) analysis. For all remaining adherent wells, Stage 3 media was aspirated and replaced with 1mL DPBS/well. Adherent cell monolayers were manually scraped with a cell scraper (Falcon #353085) and collected using a serological pipette into 15mL conical tubes. Typically, two 6-well plates were collected per 15mL conical, and up to eight 6-well plates were collected per line. The 15mL conical tubes were centrifuged for 2 minutes at 161g. The supernatant was then aspirated and discarded, and the pellets were re-suspended in 1mL DPBS by gentle trituration using a P-1000 pipette. Once resuspended, all pellets were combined in a final volume of approximately 10mL DPBS and centrifuged for 2 minutes at 161g. Again, the supernatant was aspirated and discarded. The pellet was then resuspended in 6mL DPBS using a 5mL serological pipette and aliquoted to six 1.7mL Eppendorf tubes (1mL/Eppendorf tube). The Eppendorf tubes were centrifuged for 4 minutes at 161g, and the supernatants were aspirated and discarded. Four of the Eppendorf tubes were snap frozen in an ethanol/dry ice slurry and stored at −80°C until shipment to omics centers for analysis. The remaining two pellets were re-suspended in 1mL each of CryoStor CS10 (Biolife Solutions #210102) using a P-1000 pipette (typically, 2-4 trituration’s were sufficient to resuspend the pellets) and each pellet was transferred to an individual cryovial (Thermo Scientific #5000-1020). CryoStor vials were then stored in a Mr. Frosty (Nalgene #5100-0001) at −80°C for 24 hours, at which time they were transferred to sample boxes and stored at −80°C until shipment to omics center for processing.

### Immunocytochemistry

Human iPSC-derived motor neuron cultures were plated on optical-bottom 96-well plates (Thermo, # 165305) and subsequently fixed in 4% paraformaldehyde for 15 minutes. Cells were blocked in 5% normal donkey serum with 0.1% Triton X-100 in phosphate buffered saline (PBS) and incubated with primary antibodies for 1 h at room temperature or overnight at 4°C. Cells were then rinsed and incubated in species-specific AF488, AF594, or AF647-conjugated secondary antibodies followed by Hoechst 33258 (0.5 μg/mL; Sigma) to counterstain nuclei. Cells were imaged using Molecular Devices ImageExpress Micro high-content imaging system or using Leica microscopes^7^ (**Figure 1b,c**). Primary antibodies used were as follows: mouse anti-SMI32 (Covance, 1:1,000); mouse anti-TuJ1 (β3-tubulin) (Sigma; 1:1,000-1,2,000); rabbit anti-glial fibrillary acidic protein (GFAP, Dako; 1:1000); mouse anti-Map2a/b (Sigma; 1:1000); rabbit anti-nestin (Millipore; 1:2000), Islet-1 Antibody (R&D AF1837; 1:250) and Nkx-6.1 (DSHB F55A10-s; 1:100).

### RNA-Seq

Total RNA was isolated from each sample using the Qiagen RNeasy mini kit. RNA samples for each subject (control or disease) were entered into an electronic tracking system and processed at the University of California, Irvine GHTF. RNA QC was conducted using an Agilent Bioanalyzer and Nanodrop. Our primary QC metric for RNA quality is based on RIN values (RNA Integrity Number) ranging from 0-10, 10 being the highest quality RNA. Additionally, we collected QC data on total RNA concentration and 260/280 and 260/230 ratios to evaluate any potential contamination. Only samples with RIN >8 were used for library prep and sequencing. Library prep processing was initiated with total RNA of 1µg using a Ribo-Zero Gold rRNA depletion and Truseq Stranded total RNA kit. Additionally, ERCC exFold spiked-in controls were used for further QC and downstream data analysis. Briefly, RNA was chemically fragmented and subjected to reverse transcription, end repair, phosphorylation, A-tailing, ligation of barcoded sequencing adapters, and enrichment of adapter-ligated cDNAs. RNA-Seq libraries were titrated by qPCR (Kapa), normalized according to size (Agilent Bioanalyzer 2100 High Sensitivity chip). Each cDNA library was then subjected to Illumina (HiSeq 2500) paired end (PE), 100 cycle sequencing to obtain approximately 50-65M PE reads. After sequencing fastq were subject to QC measures and reads with quality scores (>Q20) collected and analyzed using the pipeline described at http://neurolincs.org/pipelines/. Briefly, reads were mapped to the GRCh37 reference genome, QCed, and gene expression and differential expression were quantified using tools HTseq^97^ and DESeq2^98^. Normalized and transformed count data were then used for exploratory analysis and DE genes (FDR <0.1) were used for pathway, network, and gene ontology analysis. These primary data were subject to additional statistical and network-based data analyses using commercial and open-source pathway and network analysis tools, including Ingenuity Pathway Analysis (IPA), Gorilla, Cytoscape, and other tools to identify transcriptional regulators, predict epigenomic changes, and determine potential downstream pathway and cellular functional effects. Significant DEGs (FDR<0.1) were then analyzed against genes that were found to contain exonic enriched genetic variants from the WGS. The gene expression (voom normalized and transformed values) and genotype variant pairs were analyzed by fitting a linear regression model. Adjusted R^2^ and Benjamini-Hochberg adjusted p-values were calculated, significant genes were reported at (FDR<0.1). The replication cohort was carried out using the same methods.

### Proteomics

Frozen cell pellets were lysed using a combination of lysis buffer containing SDS and sonication. BCA assay was used to determine protein concentration and 125ug of each sample was used in downstream sample processing. Samples were processed following Expedeon FASP protocol^99^. Samples were digested in Trypsin/LysC (Promega) at a ratio of 40:1 to protein concentration at 37°C for 12 hrs. Samples were desalted using MCX micro-elution column (Waters) and samples were dried in speedvac and stored in −20C until resuspension with Biognosys iRT mixture for acquisition on the SCIEX 6600 over a 45-minute gradient. Samples were acquired in data-dependent acquisition (DDA) mode for library building and in data-independent acquisition (DIA) mode over 100 variable windows similar to acquisition protocols in Kirk et al. and Holewinski et al.^100, 101^. DDA files were run through Trans Proteome Pipeline (TPP) using a human canonical FASTA file (Uniprot). A consensus peptide library with decoys was generated. DDA library build principals as described in Parker et al^102^ were utilized to generate a cell specific library, which allowed for more accuracy in matching DIA data to the DDA library during OpenSWATH, as indicated by higher d-scores in PyProphet. DIA files were mapped onto this library using OpenSWATH and transition level data was compiled with a 1% FDR cutoff. Downstream summing of transition level data to peptide and protein level data was performed by MAP DIA^103^. Log2FC data was calculated by MAP DIA and filtered using a 1% FDR, 95% confidence interval and 0.6 abs(log2FC) cutoff to obtain a final list of differentially expressed proteins. For protein quantification, transitions and peptides common to more than one protein were excluded. These data have been further analyzed using commercial and open-source pathway and network analysis tools, including Ingenuity pathway analysis and GOrilla to identify upstream regulators and determine affected cellular pathways.

#### Replication Cohort

The sample processing methodology for proteomic analyses was altered to enable high-throughput automation using the Biomek i7 Liquid Handling Automated Workstation. This concomitantly reduces manual processing technical variations and thus improves long-term longitudinal aspirations inherent to the project and presently ongoing. Cell pellets were lyophilized at −55°C and were solubilized in 6M Urea, 1mM DTT in 1M NH4HCO3, pH 8 and sonicated at 70% amp, 10 sec on and 10 sec off (800R1 QSONICA) at 4°C. Sample volume is diluted by 2/3 using 100mM Tris, 4mM CaCl, pH 8. Transfer 200 ug protein, as determined by BCA assay (Pierce BCA Protein Assay Kit) to 96 well reaction plate for robotic digestion on the Biomek i7 automated workstation (Beckman Coulter). Reduction and alkylation is performed using TCEP and IAA, respectively and 2ug of β-Galactosidase is added to each sample as a digestion control. Protein sample extracts are digested with 5 ug Trypsin/LysC mix (Promega) for 4 hours at 37°C. Final sample is acidified with 10% FA and transferred onto a conditioned 96 well HLB 5mg column (Waters) for desalting. Peptides are eluted from the HLB with 50% MeCN, 0.1% FA and stored at −80°C after dried to completion on a speedvac system. Mass spectral DIA data was generated on the Triple TOF 5600 (SCIEX) and the peptide ion library data was generated on the Triple TOF 6600 (SCIEX). Data analysis was performed as described above.

### ATAC seq

We used the assay for transposase-accessible chromatin using sequencing (ATAC-Seq) to assess chromatin accessibility and identify functional regulatory sites involved in driving transcriptional changes associated with C9ORF72. ATAC-Seq detects open chromatin sites and maps transcription factor binding events in regulatory elements genome-wide, without needing any prior information about which proteins are bound. By correlating ATAC-Seq patterns with other features, such as gene expression, we are able to delineate the fine-scale architecture of the regulatory framework. Chromatin accessibility signatures were generated for each sample individually with detection of differential peaks between disease and control states to generate an initial disease-state signature.

#### Initial cohort

ATAC-seq was carried out as described^104^. Briefly, cells were lysed in cell lysis buffer (10 mM Tris-HCl, pH 7.4, 10 mM NaCl, 3 mM MgCl2, 0.1% IGEPAL CA-630, protease inhibitors) on ice for 5 min and centrifuged at 230 rcf for 5 min at 4 °C. The pellet, containing the nuclei, was re-suspended in 25 µl of 1X Tagment DNA Buffer (Illumina). 50K nuclei were subjected to transposase reaction (Nextera - Illumina) followed by DNA purification. The tagmented DNA was PCR amplified using Nextera indexing primers (Illumina) and loaded on 2% agarose gel. Nucleosome-free fragment (175-250 bp) were size selected from the gel and further amplified by PCR to obtain the final libraries. The libraries were sequenced using the Illumina HiSeq 2000 platform (single end, 50 bp). All samples passed quality control checks that included morphological evaluation of nuclei, agarose gel electrophoresis of libraries, and real-time qPCR to assess the enrichment of open-chromatin sites. The quality of the sequencing was assessed using FastQC and the reads were aligned to GRCh37 genome build using BWA. We identified open chromatin regions separately for each sample using the peak-calling software MACS2^105^ and determined differentially open sites using DEseq2 (FDR<0.1). Peaks were assigned to unique genes using the default HOMER^106^ parameters, and gene ontology analysis was performed using GOrilla^107^.

### Data Integration

We used a hierarchical strategy for data integration. We inferred transcriptional regulators from the combination of ATAC-seq and RNA-Seq data, and then searched for connections among these transcriptional regulators and those detected directly by the proteomics.

#### Inferring transcriptional regulators

Accessible chromatin regions, identified by ATAC-seq, were combined with differential gene expression data to predict transcription factors (TFs) that contribute to differences in transcriptomics profiles between C9 and controls. Specifically, we used the union of peaks detected in ALS and control samples to identify peaks proximal (+/−2.5kb) and distal (+/−50kb) to gene transcription start sites (TSS), which were further divided into those with high and low CpG content. A normalized CpG metric was used as previously described^108^. We determined the enrichment of known motifs using HOMER. The analysis was performed separately for high and low CpG content peaks near (+/−10kb or 50kb) differentially expressed genes as the foreground and corresponding regions near all known genes as the background.

### Network Analysis

We used Omics Integrator^36^ to search for previously reported protein-protein interactions that link proteins detected by mass-spectrometry and the inferred transcription factors. Taking a network approach, we represented proteins and TFs as nodes and assigned prizes to them based on their experimental significance. Specifically, protein prizes were assigned according to the fold change between C9 and control samples and prizes for TFs were assigned according to false discovery rate (see above). We mapped these proteins on a network of physical interactions in which each edge was scored for reliability based on the underlying experimental data. Our algorithm searches for disease-associated subnetworks that retain the maximum prizes while avoiding unreliable interactions which are formalized as the Prize-Collecting Steiner Forest problem. We aim to find a forest solution *F(V_F_, F_F_)* that maximizes the objective function:

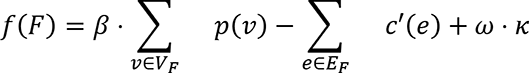

The first term is the sum of prizes included in *F*, scaled by a model parameter *β*. The second term is a cost function which serves the purpose of only including a node in *F* if the objective function is minimized. The last term allows for the inclusion of *κ* trees by introducing a root node *v*_0_ that is connected to every other node with a weight *ω*. This method not only performs feature selection by filtering out protein prizes that are expensive to connect, but also identifies “Steiner” proteins that were not detected as changing in the experiments, but are strongly implicated by the structure of the interactome. A Steiner node is typically included when its interaction neighbors are significant proteins identified from biological experiments. To avoid a bias toward proteins that have many known interactions (high-degree nodes), we impose a regularization term on edges such that the cost of an edge between nodes *a* and *b* monotonically increases with *d*_*a*_ and *d*_*b*_, the node degrees of *a* and *b*:

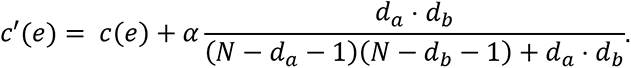

This regularization term corresponds to the probability that an edge exists between *a* and *b* given the number of nodes in the interactome, *N*, and the degrees of *a* and *b*. *c*(*e*) is the cost of the edge which is inversely related to the amount of experimental evidence supporting the physical interaction between *a* and *b* given by iRefIndex^109^. Finally, we acknowledge that the algorithm is susceptible to noise in the interactome, so we ran the experiment 100 times with randomly added noise to the interactome and chose the top 400 nodes that appeared most frequently and removed any disconnected nodes. Additionally, we assessed the specificity of the network by assigning the input prize values to random nodes in the interactome and measuring the frequency that each node appears. We repeated these experiments for a parameter grid and selected a network that 1) performed feature selection (i.e., did not include the entire input prize list), 2) was specific (as determined by the calculations using randomly assigned prizes), and 3) had a degree distribution that matched that of the input prize file. As C9ORF72 was not detected in the proteomics measurements, we forced C9ORF72 inclusion in the network by artificially assigning it a large prize. Network nodes were then sorted by subcellular location based on the Compartments database^110^ and plotted in Cytoscape.

### Drosophila Screen

An initial set of 249 genes were selected from two sources: 1) proteins from the integrated network and 2) miscellaneous genes of interest that were of interest to various group members. Out of the 249 genes, a majority (141) were selected from the integrated network. Drosophila orthologs of human DEGs were identified using DIOPT^111^, and transgenic fly lines knocking-down or overexpressing these genes downstream of UAS sites for GAL4-specific modulation were obtained from the Bloomington Drosophila Stock Center. We identified 300 fly genes corresponding to these 249 human genes (or 284 including paralogues) and conducted 334 total fly experiments (several fly genes were tested with different knockouts). Of these 334 total experiments, 7 exhibited lethal phenotype, 7 were not true modifiers as determined by GMR GAL4 score, and 1 was both. Filtering these out, we had 321 remaining fly experiments representing 288 fly genes and 242 human genes. These modifiers were crossed to flies overexpressing the hexanucleotide repeat expansion (HRE) in the eye [GMR Gal4; UAS-(G4C2)30/CyO]. Progeny co-expressing both the HRE and putative modifier were collected within 24 hours of eclosion and aged at 25°C and compared to control flies of the same genetic background. A relative modification index, ranging from −4 to +4, was used to assess eye degeneration where −4 represented complete rescue and +4 represented no eye^112^. A score of 0 represents no effect of the tested modifier. Ommatidial structure, interommatidial bristles, necrosis, loss of pigmentation, and overall morphology of the eye were assessed during scoring. Only female flies were scored due to male flies displaying a higher degree of variability. All experimental modifiers were tested with 3 biological replicates with their eye degeneration scores averaged. If a fly cross failed to eclose, the subsequent score was marked ‘lethal’. Selected strong enhancers and suppressors were retested with GMR Gal4; UAS-(G4C2)30/CyO as well as GMR Gal4 alone, at both 25°C and 29°C. At 15 days, representative female eyes were imaged using a Nikon SMZ1500 stereomicroscope and Lumenera INFINITY3-6UR 3.0 Megapixel camera and analyzed with Image-Pro Insight v9.

In some cases, a human candidate gene had multiple fly orthologs. For each human gene, a “weighted eye score” was calculated by taking the average of all corresponding fly orthologs, weighted by the ortholog scores as determined by the DRSC Integrative Ortholog Prediction Tool (https://www.flyrnai.org/cgi-bin/DRSC_orthologs.pl). Note that only moderate and high ranking orthologs were considered.

### Drosophila Network Analysis

We categorized the genes that were tested in the Drosophila model into three groups: causal, compensatory and non-contributory. For example, we reasoned that genes that were significantly upregulated in ALS and whose knockdown in fly suppressed or enhanced eye degeneration were likely causal or compensatory genes, respectively. Similarly, those that were significantly downregulated in ALS and were enhancers or suppressors of eye degenerations were likely causal or compensatory, respectively. Genes whose knockdown in the fly model showed little to no effect on eye degeneration were categorized as non-contributory.

Next, we used previously annotated directed interactions that were pulled from the ReactomeFiViz and KEGG databases^113, 114^. The resulting directed network was composed of 9,336 nodes connected by 166,907 directed edges. For any two proteins that were labeled as either causal or compensatory, we identified all directed paths of length at most 2. Next, we only considered paths that were concordant with our data by not allowing paths:

1. to contain genes that are *not* expressed in iMNs. This was defined by taking the top 70% of expressed gene transcripts across all 7 iMNs lines.
2. whose predicted effect on protein activity is discordant with measured protein expression. For instance, if A->B, but A is up in ALS and B is down in ALS, this edge is excluded from further analysis. Direction of interaction (activating or inhibiting) was extracted from ReactomeFiViz and KEGG databases^113, 114^ to determine the predicted effect.

### Statistical Analysis

#### Immunostaining

The boxplots shown in Figure 1C are average results from quantified images of the respective immunostains in Figure 1B. The healthy control donors (CTR) comprised of n = 3 independent iPSC lines, while the C9-ALS comprised of n = 4 C9ORF72 repeat expansion donor iPSC lines. Total cells were quantified by nuclear staining with Hoechst 33258 in n = 9 sites across a well and percent positive cells for respective marker were calculated for each site. Average positive marker expression was then calculated for each well. Each marker immunostain was performed across independent well 3 times and respective average percent positive cells were obtained for each iPSC lines. All statistical analyses for percent SMI32, TuJ1, Map2a/b, GFAP and Nestin levels were performed using unpaired t test and the differences between CTR and C9-ALS groups were insignificant. Error bars represent SEM. *RNA-seq*: Generalized linear models were used with negative binomial distribution to estimate fold change between ALS and controls samples for each gene. Wald test was performed for hypothesis testing, which is a one-sided test. Sample size n was 3 and 4 respectively for control and ALS in the first cohort and 4 and 6, respectively in the second cohort. Significant DEGs and enrichment terms were chosen based on a 10% FDR. *Proteomics*: Throughout Trans Proteome Pipeline (TPP) and OpenSWATH, a 1% FDR cutoff was employed in identification of transitions/peptides and in OpenSWATH matching to the peptide library. MAP DIA^103^ was used on MS2 normalized transition level data obtained from OpenSWATH. Transitions falling outside of 2 standard deviations were filtered out. Additional correlation filter of 0.2 was used to filter out any residual outliers. Intensities of the remaining transitions were summed for peptide, and then protein level quantification. Differential expression analysis of designated groups was performed by MAP DIA using analysis based on a Bayesian latent variable model with Markov random field prior. Output for differential expression included log2FC, confidence score, FDR and log(OddsofDifferentialExpression). Log2 fold changes were deemed significant if they had FDR at 1% or lower, a confidence score of .95 or above, a positive log(oddsDE) and an abs(log2FC) of .6 or above. For IPA analysis, the 924 differentially expressed proteins and their corresponding log2FC values were used, with analysis settings for reference set: Ingenuity Knowledge Bases, direct relationships, using all data sources, experimentally observed interactions and filtered for human genes in primary tissues and human cell lines. For pathway enrichment analysis, GOrilla^107^ was used. The DIA filtered list of 3742 proteins was used as the background list for analysis of target sets. A p-value threshold of 10^-3^ was used to determine enriched GO Biological Process terms. *ATAC-seq*: Differentially open sites were called using the DESEQ2 pipeline with FDR <= 0.1. Data integration: All GO enrichments were performed using a one-sided hypergeometric test implemented by GOrilla. Motif enrichments were calculated via HOMER which searches for de novo motif matches that are enriched in a set of foreground sequences relative to a given set of background sequences using a one-sided hypergeometric test. Enrichment of ALS-associated genes was calculated using a one-sided hypergeometric implemented using the hypergeometric module in Scipy v0.14. Enrichments of genes between -omics assays were also calculated using a one-sided hypergeometric test implemented using the hypergeometric module in Scipy v0.14. For each pair of assays, the background was the set of genes that was detected in both assays. *Drosophila eye screen*: flies were aged to 15 days after eclosion. 3 biological replicates were carried out per cross. 15 females flies were scored per cross. The average score of these 15 flies was taken as the average for one biological replicate. The average of all 3 biological replicates rounded to the nearest 0.5 of a point was used for the final rounded rough eye score.

### Data and Code Availability

All data and code are available from the corresponding authors upon reasonable request.

#### Accession Codes

ATAC-seq and RNA-seq raw data is available in dbGAP (https://www.ncbi.nlm.nih.gov/gap) and processed data is available through the LINCS portal (http://lincsportal.ccs.miami.edu/dcic-portal/). Proteomics raw data is available in the CHORUS database (https://chorusproject.org/pages/dashboard.html#/search/Neurolinc/projects) and processed data available through the LINCS portal (http://lincsportal.ccs.miami.edu/dcic-portal/). Replication cohort data submission is pending.

## Supplemental Figure Legends

**Figure S1:** Schematic for generation of (A) iPSC-derived motor neurons precursor spheres (iMPS) and (B) iMPS-derived motor neurons (iMNs), and media components for each stage.

**Figure S2:** (A) Schematic for generation of the replication cohort of CTR and C9-ALS iPSCs into direct iPSC-derived motor neurons (diMNs) cultures using a rapid 3 stage protocol used by NeuroLINCS for transcriptomics, proteomics and ATAC-seq assays. (B) Violin plots quantifying levels of NEFH (SMI32). Islet1, Nkx6.1, and Tuj1 in control and C9-ALS diMNs cultures. Percent ISLET1 positive cell count in CTR vs C9-ALS groups as statistically significant (***).Two-tailed P value = 0.0009; Unpaired t test with Welch’s correction. CTR n=7 and C9-ALS n=6.

**Fig. S3:** Representative images of diMNs from 7 control and 6 C9-ALS iPSCs from the replication cohort largely showing distribution of neural cell populations marked by SMI32 (NEFH), Islet1, Nkx6.1, and Tuj1 (TUBB3) in control and C9-ALS diMNs cultures. For each set of stains and for every cell line differentiated into diMNs, there is dotted box in the main image which shows the region that is magnified in the adjacent image. All scale bars are 100 µm.

**Figure S4:** (A) G-band karyotypes depict normal cytogenetic profiles in the 3 control iPSC lines and the 4 C9-ALS iPSC lines used in this study. (p, passage at which cells were harvested). (B) G-band karyotypes depict normal cytogenetic profiles in the replication cohort of the additional 7 C9-ALS and the 6 control iPSC lines used in this study. (P, passage at which cells were harvested).

**Figure S5: (A**) DNA Fingerprinting human 9 species-specific short-tandem repeat (STR) marker profiling confirms the that the reprogrammed iPSCs and the differentiated iMNs used for ‘OMICS assays in this study match the parental donor fibroblasts. * N/A (not applicable) as the parent fibroblast line was not available for comparison purposes. The genetic profile for the sample was compared to the cell line genetic profiles available in the DSMZ STR database and to all previously submitted profiles in the Cedars-Sinai iPSC Core. The profiles were found to be unique and did not match to any previously submitted profiles. The genetic profile established for this sample can be used for future comparisons for this cell line. (B) DNA Fingerprinting human 9 species-specific short-tandem repeat (STR) marker profiling confirms that the replication cohort of the reprogrammed iPSCs and the differentiated diMNs used for ‘OMICS assays in this study match the parental donor PBMCs and iPSCs. * N/A (not applicable) as the parent PBMCs were not available for comparison purposes. The genetic profile for the sample was compared to the cell line genetic profiles available in the DSMZ STR database and to all previously submitted profiles in the Cedars-Sinai iPSC Core. The profiles were found to be unique and did not match to any previously submitted profiles. The genetic profile established for this sample can be used for future comparisons for this cell line.

**Figure S6:** (A) Volcano plots of log2 fold change and −log2(adjusted pvalue) from RNAseq data for first (oldiMN) and second (DE4) cohorts. (B) MA plots of log2 fold change and mean normalized counts from RNAseq data for first (oldiMN) and second (DE4) cohorts. (C) Cell Type-Specific Analysis of 828 DEGs from the iMNs. Revealing an enrichment for cortical and motor neurons. (D) NRG1 subnetwork showing upregulated target genes (Red) from C9 vs Control iMN and predicted activation of upstream regulators (Orange). (E) Fold change levels of MMPs and associated substrates in ALS-C9 vs control iMNs. (F) Percent alternative splicing types in C9 iMNs. (G-I) GO enrichment analysis of significant differentially spliced genes.

**Figure S7:** Proteomics Data QC. A) Mass spectrometry (MS) runs result in 3844 unique hits, based on 23,436 peptides. Shared hits are indicative of peptides that mapped to multiple proteins; these were not used in further data analysis. B) Total ion current distribution for each file shows similar levels of MS1 and MS2 TIC. C) Log2 Protein Intensity distribution of MS2 normalized protein data shows the spread of protein intensities for each MS run. There are no major differences between sample distribution and no major difference in intensities between control and disease samples. D) ALS and Control normalized protein intensity show a high correlation, with R = .9660. This shows that both ALS and Control sample differentiations yielded similar samples. E) Correlation between overlapping differentially expressed proteins and genes is high. F) Principal component analysis shows separation of ALS and Control in PC1 when PC1 and PC4 are mapped. G) IPA analysis shows predicted inhibition of RNA processing and splicing.

**Figure S8:** (A) A histogram of differential and consensus ATAC-seq peaks’ distance to their nearest genes show that differential peaks lie further away from genes on average. (B) Consensus (left) and differential (right) peaks were mapped to Gencode annotations. A smaller proportion of differential peaks lie in promoters than consensus peaks. (C) Roughly 6% of all consensus peaks are open in ALS, and another 6% are open in control. The two sets of tracks on the right are examples from both of these categories. (D) Each differential peak was assigned to the nearest gene within 50kb and the number of ALS and CTR peaks were counted for each gene. The heatmap intensity indicates the number of genes with a given combination of ALS and CTR peak counts.

**Figure S9:** eQTL: Boxplots showing each significant genotype-gene expression comparison of genes also found in our integrated network. Y-axis: gene expression, X-axis: genotype. Genotype (Red = Ref, Green = Het, Blue = Homo). Dots are individual replicates of each sample in study (ALS= Red, Control = Blue).

**Figure S10:** eQTL: Boxplots showing all genotype-gene expression comparisons for known brain eQTLs found in our WGS and also found in our significant DEGs. Y-axis: gene expression, X-axis: genotype. Genotype (Red = Ref, Green = Het, Blue = Homo). Dots are individual replicates of each sample in study (ALS= Red, Control = Blue). Variant rsID are listed below each plot.

**Figure S11:** Top GO enrichments for each assay.

**Figure S12:** Subnetwork of disease network from Figure 4C explores possible effects of decreased Sumoylation in C9-ALS lines.

**Figure S13:** Density and histogram plots from randomization tests. (A) Density plot of −log(p values) showing significance of overlap between 100 network randomizations and DEGs from ALS vs CTR postmortem cervical spine (FDR < 0.1), calculated by Fisher’s Exact test. Dashed orange line marks −log(p value) of true integrated network (9.18E-05) compared to the postmortem DEGs. (B) Density plot of number of overlapping genes between 100 network randomizations and DEGs from ALS vs CTR postmortem cervical spine (FDR < 0.1). Dashed orange line marks the number of overlapping genes from the true integrated network (81 genes) compared to the postmortem DEGs. (C) Histogram and distribution of the numbers of DEGs found using 1000 randomized permutations. Red line marks value generated using actual sample labels. (D) Histogram and distribution of overlap between 1000 randomized DEG lists and our ECM subnet. Red line marks value generated using actual sample labels.

**Figure S14:** Density plot showing change in eye phenotype and change in protein expression for each fly perturbation shows no correlation.

**Figure S15:** Comparison of proteins from the integrated network (Figure 5A) between two sets of cultured motor neuron experiments. The horizontal and vertical components of the arrows indicate protein fold changes (ALS/CTR) between the original and validation experiment, respectively. Arrows colored black indicate proteins whose fold changes were in the same directions between experiments. Arrows colored red indicate proteins whose fold changes were in different directions between experiments.

## Supplemental Tables

**Tables 1-4: Reagents for Cell Differentiations**

**Table S5**: Clinical and reprogramming details for the ALS-C9 and control iPSC lines used in the NeuroLINCS study. W: White; AA: African American; M: Male; F: Female; CS: Cedars-Sinai; HD: Huntington Disease.

**Table S6**: Summary Table of all variants in iPSC lines from healthy volunteers and people with ALS due to C9ORF72 mutation. The total number of variants that are exonic functional are reported. These are nonsynonymous variants which include missense, splicing. frameshift, non-frameshift, stop-gain and start loss variants only. For regulatory variants, we filtered for variants that are in intergenic and regulatory regions. We report the variant as found next to the closest gene, these will be either in the 5’ or 3’ UTR, intronic, upstream and downstream up to 4 KBs from the start and stop of a gene.

**Table S7**: Summary Table of all ALS variants in the control and C9ORF72 lines. ALS specific variants were found in one of the CS83iCTR-33n1 and 2 of the C9ORF72 lines.

**Table S8**: Summary table of the exonic functional variants within all lines the variants that are enriched in either the cases or control.

**Table S9**: List of genes analyzed for statistically significant differences from RNAseq. See Excel sheet Table S6.

**Table S10**: Summary of Proteomics. See Excel Sheet Table S7 and corresponding tabs.

**Table S11**: Summary table of the exonic functional variants within all lines the variants that are enriched in either the cases or controls.

**Table S12**: Genes used for linear regression analysis of gene variant and expression comparisons.

**Table S13**: List of ALS associated genes from text mining sources and experimental sources.

**Table S14**: Summary table of RNAi lines used and full G4C2 fly screen results.

**Table S15**: Proteomics fold changes (ALS/CTR) and false discovery rates (FDR) from the original and validation experiments for proteins in the fly network (Figure 5).

## Notes

### Competing Interest Statement

The authors have declared no competing interest.

