## Supplemental Figures for "An integrated multi-omic analysis of iPSC-derived motor neurons from C9ORF72 ALS patients"

Supplementary Figure 1

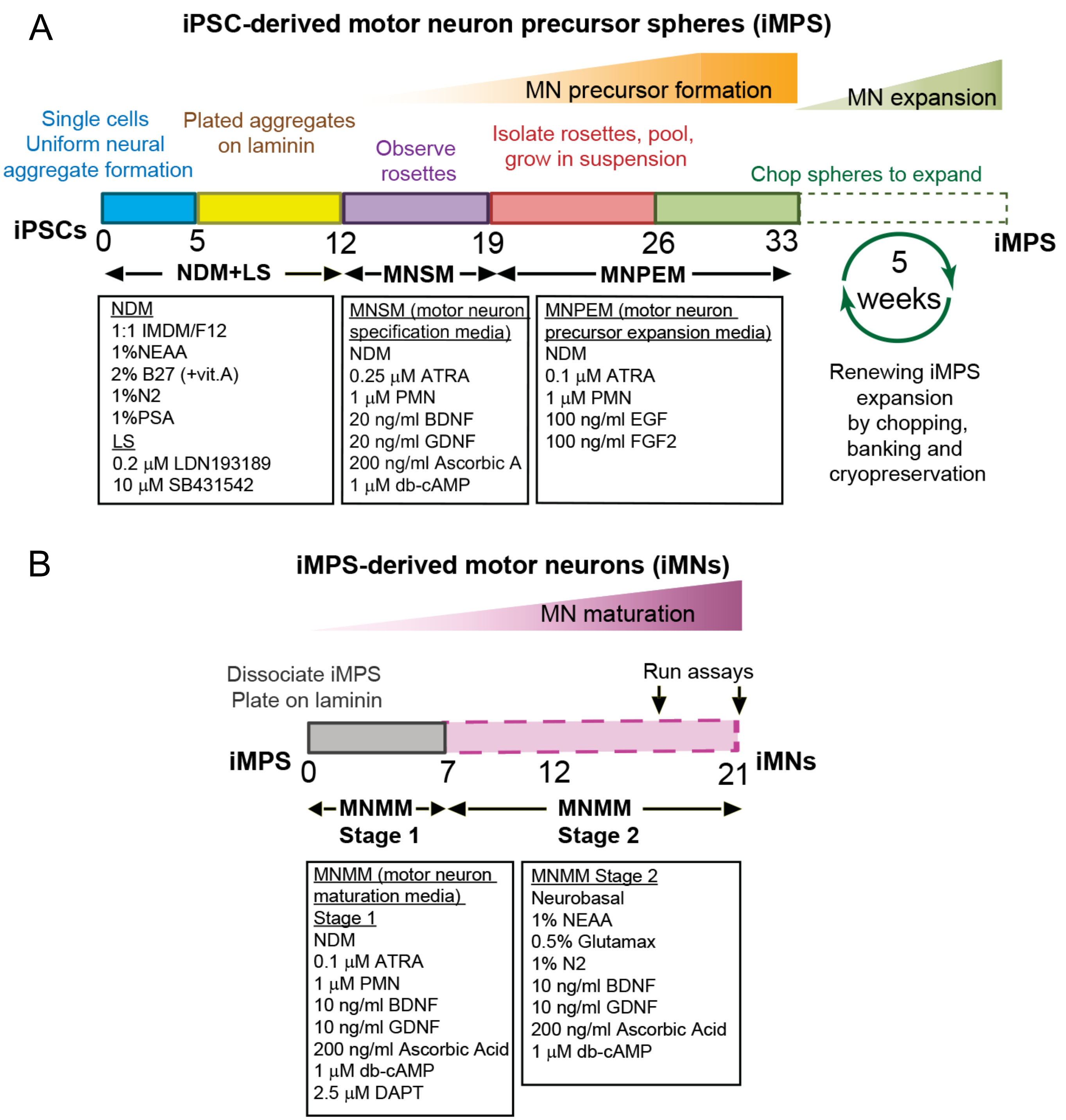

Supplementary Figure 2

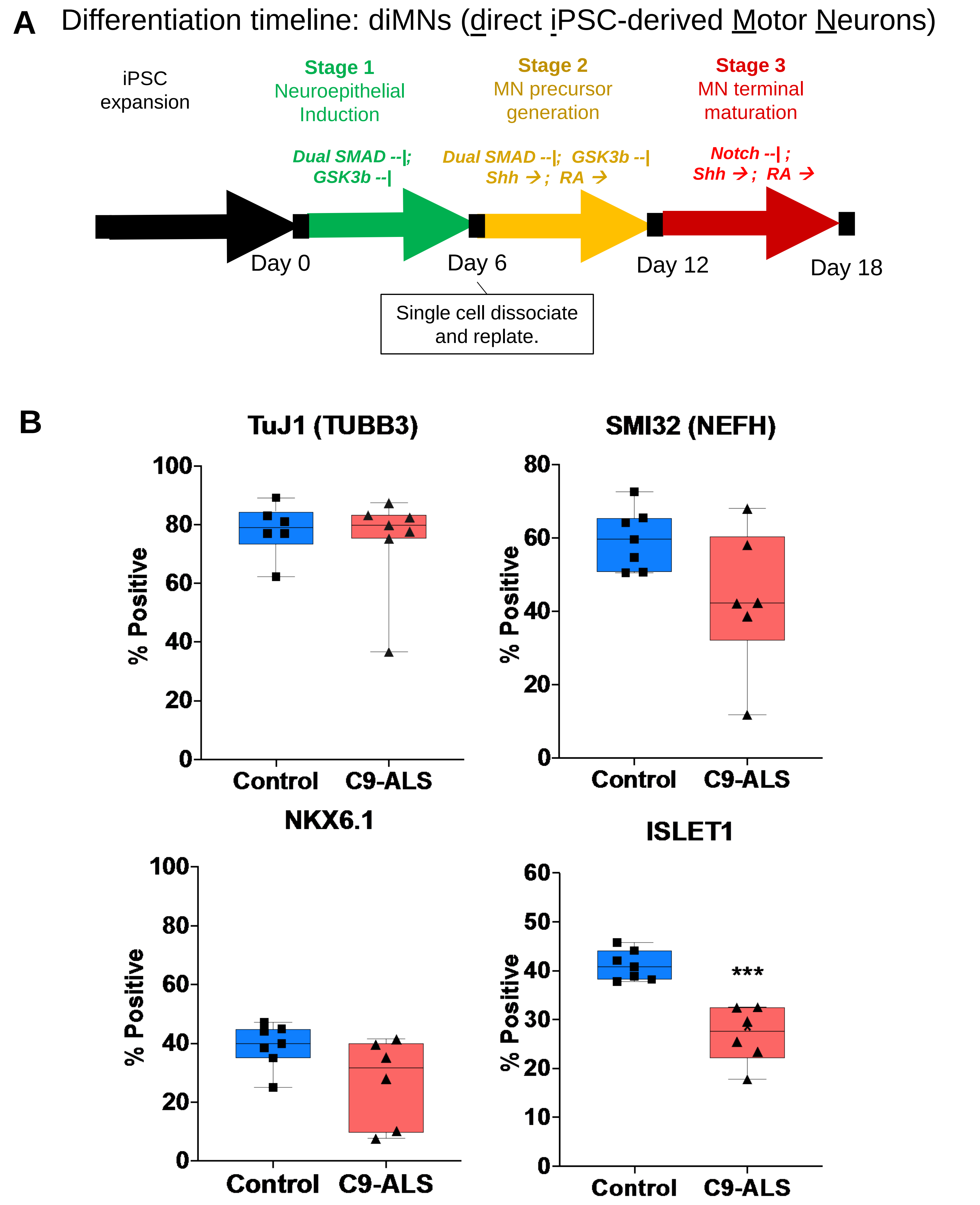

Supplementary Figure 3

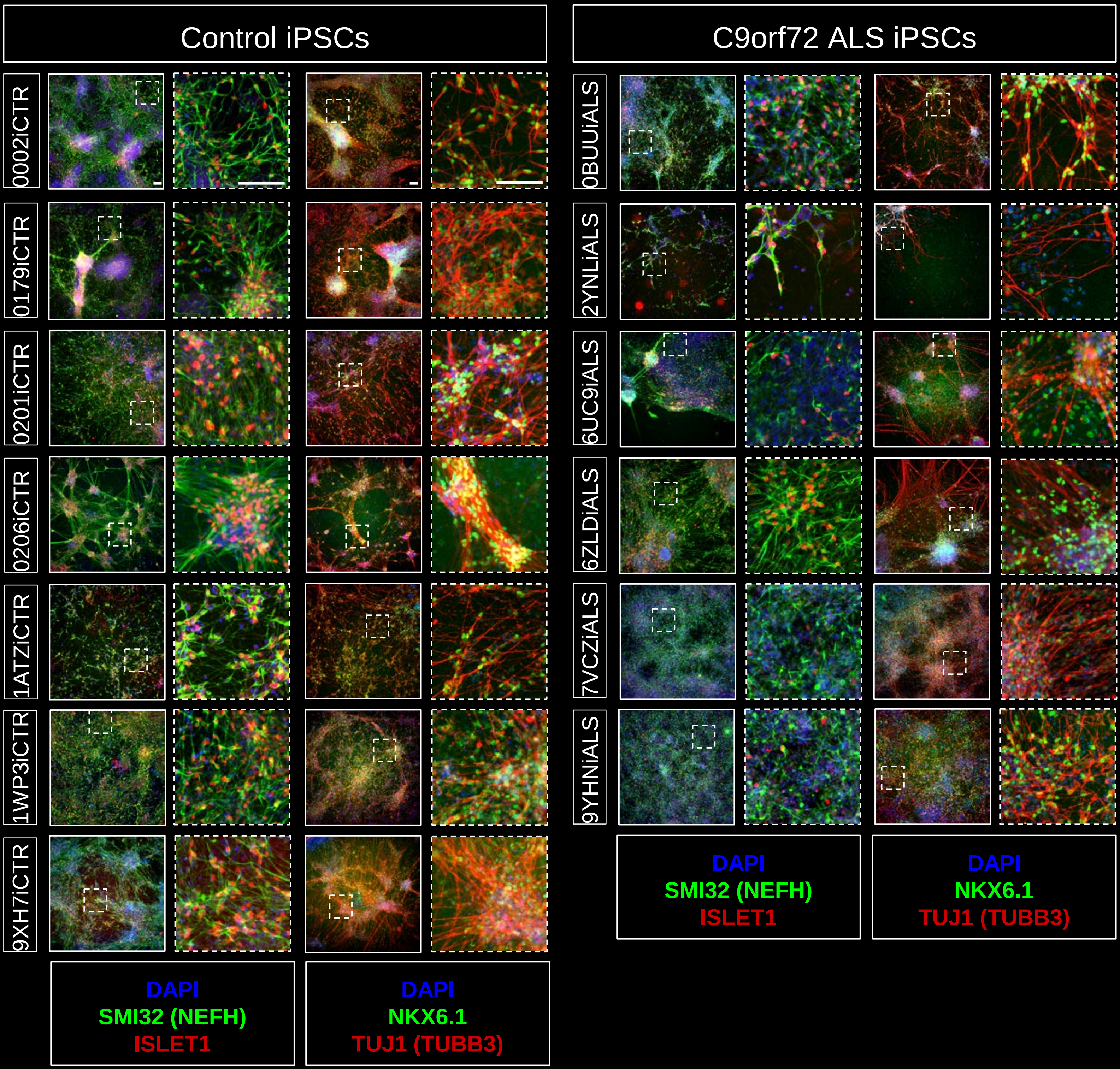

Supplementary Figure 4A G-band Karyotypes of iPSCs

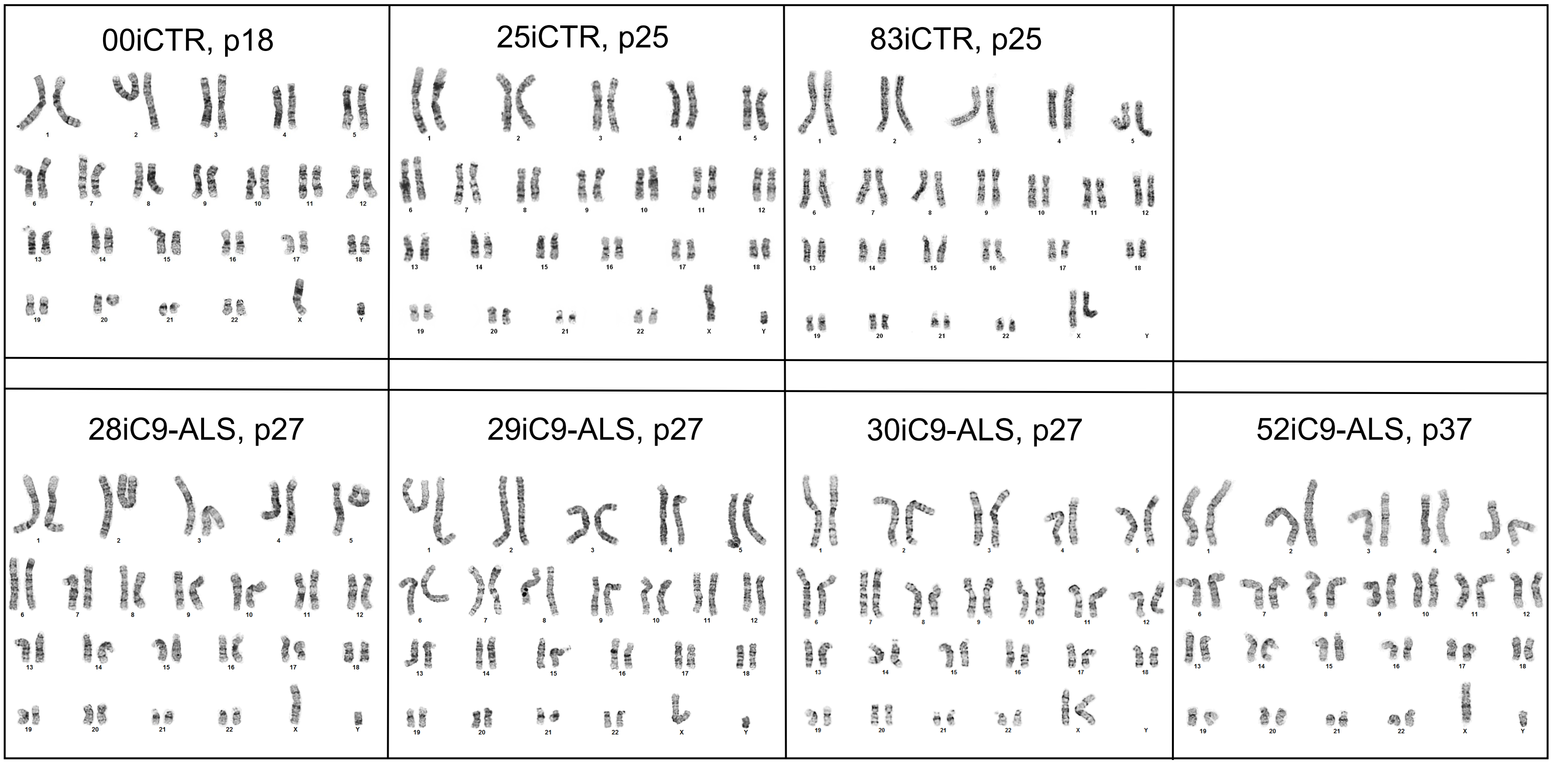

Supplementary Figure 4B

| | | | |
|---|---|---|---|
| CS0BUUiALS-n3, P19 | CS2YNLiALS-n1, P18 | CS6UC9iALS-n1, P19 | CS6ZLDiALS-n1, P18 |
| 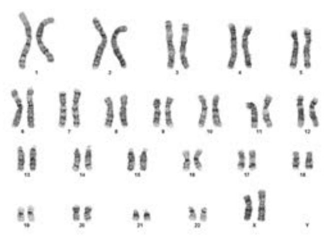 | 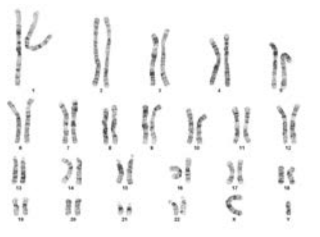 | 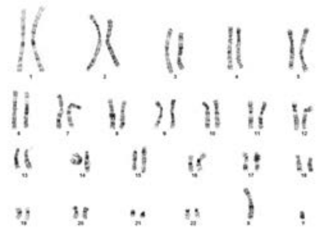 | 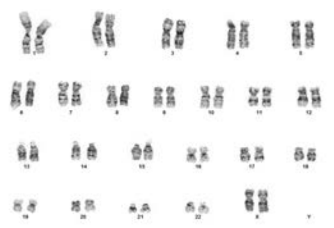 |
| CS7VCZiALS-n3, P19 | CS9YHNiALS-n1, P18 | | |
| 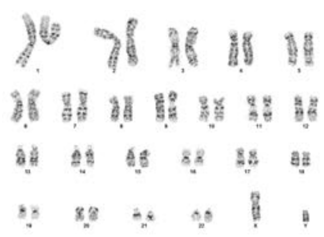 | 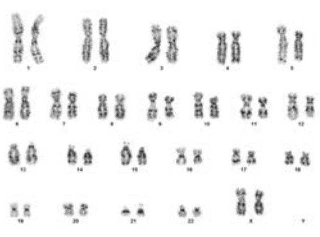 | | |
| CS0201iCTR-n4, P19 | CS0206iCTR-n5, P20 | CS1ATZiCTR-n2, P18 | CS1WP3iCTR-n8, P20 |
| 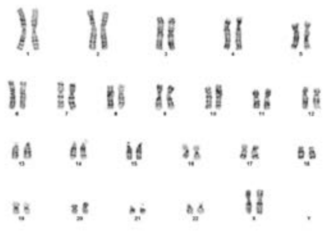 | 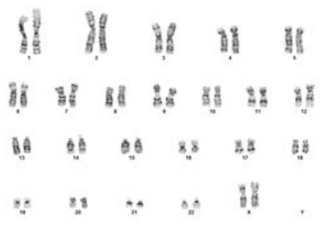 | 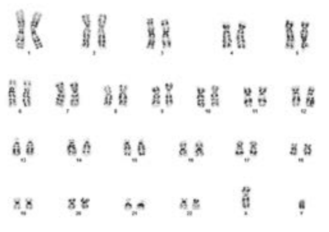 | 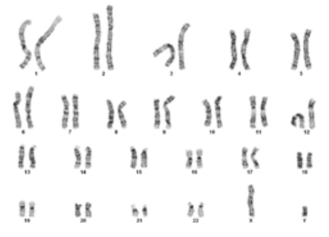 |
| CS9XH7iCTR-n4, P21 | CS0002iCTR-n1, P21 | CS0179iCTR-n1, P22 | |
| 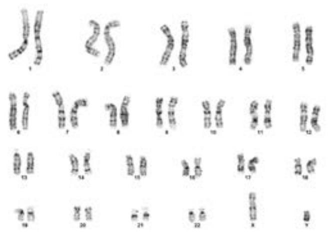 | 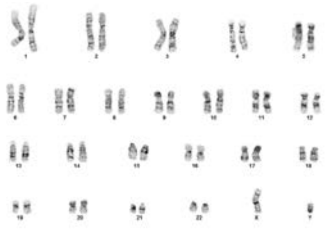 | 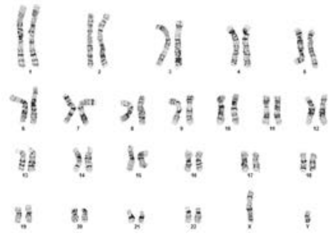 | |

Supplementary Figure 5A

### STR of iPSC Lines and iMNs

| Cell Line | Cell Type | AMEL | CSF1PO | D13S317 | D16S539 | D5S818 | D7S820 | TH01 | TPOX | vWA | % Match |
|---|---|---|---|---|---|---|---|---|---|---|---|
| | | Marker Name | | | | | | | | | |
| GM05400 | Parent Line | X, Y | 8, 13 | 11, 12 | 11, 12 | 8, 12 | 8, 9 | 6, 8 | 9, 11 | 16, 17 | - |
| 00iCTR | iPSC | X, Y | 8, 13 | 11, 12 | 11, 12 | 8, 12 | 8, 9 | 6, 8 | 9, 11 | 16, 17 | 100% |
| 00iCTR | iMN | X, Y | 8, 13 | 11, 12 | 11, 12 | 8, 12 | 8, 9 | 6, 8 | 9, 11 | 16, 17 | 100% |
| ND30625 | Parent Line | X, Y | 12 | 12, 13 | 11, 12 | 11, 12 | 9, 10 | 9.3 | 8 | 18 | -% |
| 25iCTR | iPSC | X, Y | 12 | 12, 13 | 11, 12 | 11, 12 | 9, 10 | 9.3 | 8 | 18 | 100% |
| 25iCTR | iMN | X, Y | 12 | 12, 13 | 11, 12 | 11, 12 | 9, 10 | 9.3 | 8 | 18 | 100% |
| GM02183 | Parent Line | X | 12 | 14 | 11 | 11, 12 | 8, 12 | 9, 9.3 | 8 | 16, 19 | - |
| 83iCTR | iPSC | X | 12 | 14 | 11 | 11, 12 | 8, 12 | 9, 9.3 | 8 | 16, 19 | 100% |
| 83iCTR | iMN | X | 12 | 14 | 11 | 11, 12 | 8, 12 | 9, 9.3 | 8 | 16, 19 | 100% |
| F09128 * | Parent Line | N/A | N/A | N/A | N/A | N/A | N/A | N/A | N/A | N/A | - |
| 28iAL | iPSC | X, Y | 12 | 11, 12 | 10, 12 | 11, 12 | 10, 12 | 6, 8 | 8, 9 | 16, 17 | - |
| 28iALS | iMN | X, Y | 12 | 11, 12 | 10, 12 | 11, 12 | 10, 12 | 6, 8 | 8, 9 | 16, 17 | 100% |
| F09229 | Parent Line | X, Y | 11, 13 | 11, 14 | 11, 12 | 11, 12 | 11, 12 | 7, 9 | 8, 9 | 15, 17 | - |
| 29iALS | iPSC | X, Y | 11, 13 | 11, 14 | 11, 12 | 11, 12 | 11, 12 | 7, 9 | 8, 9 | 15, 17 | 100% |
| 29iALS | iMN | X, Y | 11, 13 | 11, 14 | 11, 12 | 11, 12 | 11, 12 | 7, 9 | 8, 9 | 15, 17 | 100% |
| F10-330 * | Parent Line | N/A | N/A | N/A | N/A | N/A | N/A | N/A | N/A | N/A | - |
| 30iALS | iPSC | X | 12 | 9, 14 | 11, 12 | 13 | 10 | 7, 9.3 | 8 | 16, 17 | - |
| 30iALS | iMN | X | 12 | 9, 14 | 11, 12 | 13 | 10 | 7, 9.3 | 8 | 16, 17 | 100% |
| F09152 | Parent Line | X, Y | 10, 11 | 8 | 9, 11 | 12 | 9, 10 | 8, 9.3 | 8, 11 | 16, 17 | - |
| 52iALS | iPSC | X, Y | 10, 11 | 8 | 9, 11 | 12 | 9, 10 | 8, 9.3 | 8, 11 | 16, 17 | 100% |
| 52iALS | iMN | X, Y | 10, 11 | 8 | 9, 11 | 12 | 9, 10 | 8, 9.3 | 8, 11 | 16, 17 | 100% |

### Supplementary Figure 5B

| Cell Line | Cell Type | AMEL | CSF1PO | D13S317 | D16S539 | D5S818 | D7S820 | TH01 | TPOX | vWA | % Match |
|---|---|---|---|---|---|---|---|---|---|---|---|
| NEUEM720BUU | PBMC | X | 10 | 11,12 | 12 | 12,13 | 11,12 | 6 | 8 | 15,18 | - |
| CS0BUUiALS | iPSC | X | 10 | 11,12 | 12 | 12,13 | 11,12 | 6 | 8 | 15,18 | 100% |
| CS0BUUiALS | diMNs | X | 10 | 11,12 | 12 | 12,13 | 11,12 | 6 | 8 | 15,18 | 100% |
| NEUVX902YNL | PBMC | - | - | - | - | - | - | - | - | - | - |
| CS2YNLiALS | iPSC | X,Y | 12,13 | 12,14 | 10,11 | 11 | 8,11 | 6 | 8 | 16,18 | - |
| CS2YNLiALS | diMNs | X,Y | 12,13 | 12,14 | 10,11 | 11 | 8,11 | 6 | 8 | 16,18 | 100% |
| NEUUL256UC9 | PBMC | X,Y | 12 | 8 | 13 | 12 | 10,12 | 6,9.3 | 8 | 18,20 | - |
| CS6UC9iALS | iPSC | X,Y | 12 | 8 | 13 | 12 | 10,12 | 6,9.3 | 8 | 18,20 | 100% |
| CS6UC9iALS | diMNs | X,Y | 12 | 8 | 13 | 12 | 10,12 | 6,9.3 | 8 | 18,20 | 100% |
| NEUPK546ZLD | PBMC | X | 12,13 | 8,12 | 11,12 | 10,12 | 12 | 9.3 | 11 | 16,18 | - |
| CS6ZLDiALS | iPSC | X | 12,13 | 8,12 | 11,12 | 10,12 | 12 | 9.3 | 11 | 16,18 | 100% |
| CS6ZLDiALS | diMNs | X | 12,13 | 8,12 | 11,12 | 10,12 | 12 | 9.3 | 11 | 16,18 | 100% |
| NEUFV237VCZ | PBMC | X,Y | 10 | 11,12 | 11 | 11 | 9,10 | 9.3 | 8,11 | 19 | - |
| CS7VCZiALS | iPSC | X,Y | 10 | 11,12 | 11 | 11 | 9,10 | 9.3 | 8,11 | 19 | 100% |
| CS7VCZiALS | diMNs | X,Y | 10 | 11,12 | 11 | 11 | 9,10 | 9.3 | 8,11 | 19 | 100% |
| NEUDT709YHN | PBMC | X | 12 | 8,12 | 11,13 | 11,12 | 9,10 | 7,9 | 10,12 | 17,20 | - |
| CS9YHNiALS | iPSC | X | 12 | 8,12 | 11,13 | 11,12 | 9,10 | 7,9 | 10,12 | 17,20 | 100% |
| CS9YHNiALS | diMNs | X | 12 | 8,12 | 11,13 | 11,12 | 9,10 | 7,9 | 10,12 | 17,20 | 100% |
| W15-C201 | PBMC | X | 10,11 | 11,12 | 9,11 | 11,14 | 8,11 | 9,9.3 | 8 | 15,18 | - |
| CS0201iCTR | iPSC | X | 10,11 | 11,12 | 9,11 | 11,14 | 8,11 | 9,9.3 | 8 | 15,18 | 100% |
| CS0201iCTR | diMNs | X | 10,11 | 11,12 | 9,11 | 11,14 | 8,11 | 9,9.3 | 8 | 15,18 | 100% |
| W15-C206 | PBMC | X | 10,12 | 12 | 11,13 | 11,12 | 9,10 | 9,9.3 | 8,11 | 14,17 | - |
| CS0206iCTR | iPSC | X | 10,12 | 12 | 11,13 | 11,12 | 9,10 | 9,9.3 | 8,11 | 14,17 | 100% |
| CS0206iCTR | diMNs | X | 10,12 | 12 | 11,13 | 11,12 | 9,10 | 9,9.3 | 8,11 | 14,17 | 100% |
| NEUMN061ATZ | PBMC | X,Y | 12,13 | 12 | 11,12 | 10,13 | 10,11 | 9, 9.3 | 8 | 14,19 | - |
| CS1ATZiCTR | iPSC | X,Y | 12,13 | 12 | 11,12 | 10,13 | 10,11 | 9, 9.3 | 8 | 14,19 | 100% |
| CS1ATZiCTR | diMNs | X,Y | 12,13 | 12 | 11,12 | 10,13 | 10,11 | 9, 9.3 | 8 | 14,19 | 100% |
| NEUVW301WP3 | PBMC | X,Y | 10,12 | 8,12 | 9,11 | 12 | 8,9 | 6,7 | 9,11 | 17,18 | - |
| CS1WP3iCTR | iPSC | X,Y | 10,12 | 8,12 | 9,11 | 12 | 8,9 | 6,7 | 9,11 | 17,18 | 100% |
| CS1WP3iCTR | diMNs | X,Y | 10,12 | 8,12 | 9,11 | 12 | 8,9 | 6,7 | 9,11 | 17,18 | 100% |
| NEUPW469XH7 | PBMC | X,Y | 11,12 | 12,14 | 11,12 | 12,13 | 8,11 | 9,9.3 | 8 | 15,18 | - |
| CS9XH7iCTR | iPSC | X,Y | 11,12 | 12,14 | 11,12 | 12,13 | 8,11 | 9,9.3 | 8 | 15,18 | 100% |
| CS9XH7iCTR | diMNs | X,Y | 11,12 | 12,14 | 11,12 | 12,13 | 8,11 | 9,9.3 | 8 | 15,18 | 100% |
| CS-002 | PBMC | X,Y | 12 | 9,11 | 9,10 | 11,12 | 10,11 | 6,9.3 | 8,11 | 14,16 | - |
| CS0002iCTR | iPSC | X,Y | 12 | 9,11 | 9,10 | 11,12 | 10,11 | 6,9.3 | 8,11 | 14,16 | 100% |
| CS0002iCTR | diMNs | X,Y | 12 | 9,11 | 9,10 | 11,12 | 10,11 | 6,9.3 | 8,11 | 14,16 | 100% |
| W14-C179 | PBMC | X,Y | 10,12 | 11 | 11 | 10,11 | 10,13 | 6,9.3 | 8 | 17,19 | - |
| CS0179iCTR | iPSC | X,Y | 10,12 | 11 | 11 | 10,11 | 10,13 | 6,9.3 | 8 | 17,19 | 100% |
| CS0179iCTR | diMNs | X,Y | 10,12 | 11 | 11 | 10,11 | 10,13 | 6,9.3 | 8 | 17,19 | 100% |

Supplementary Figure 6

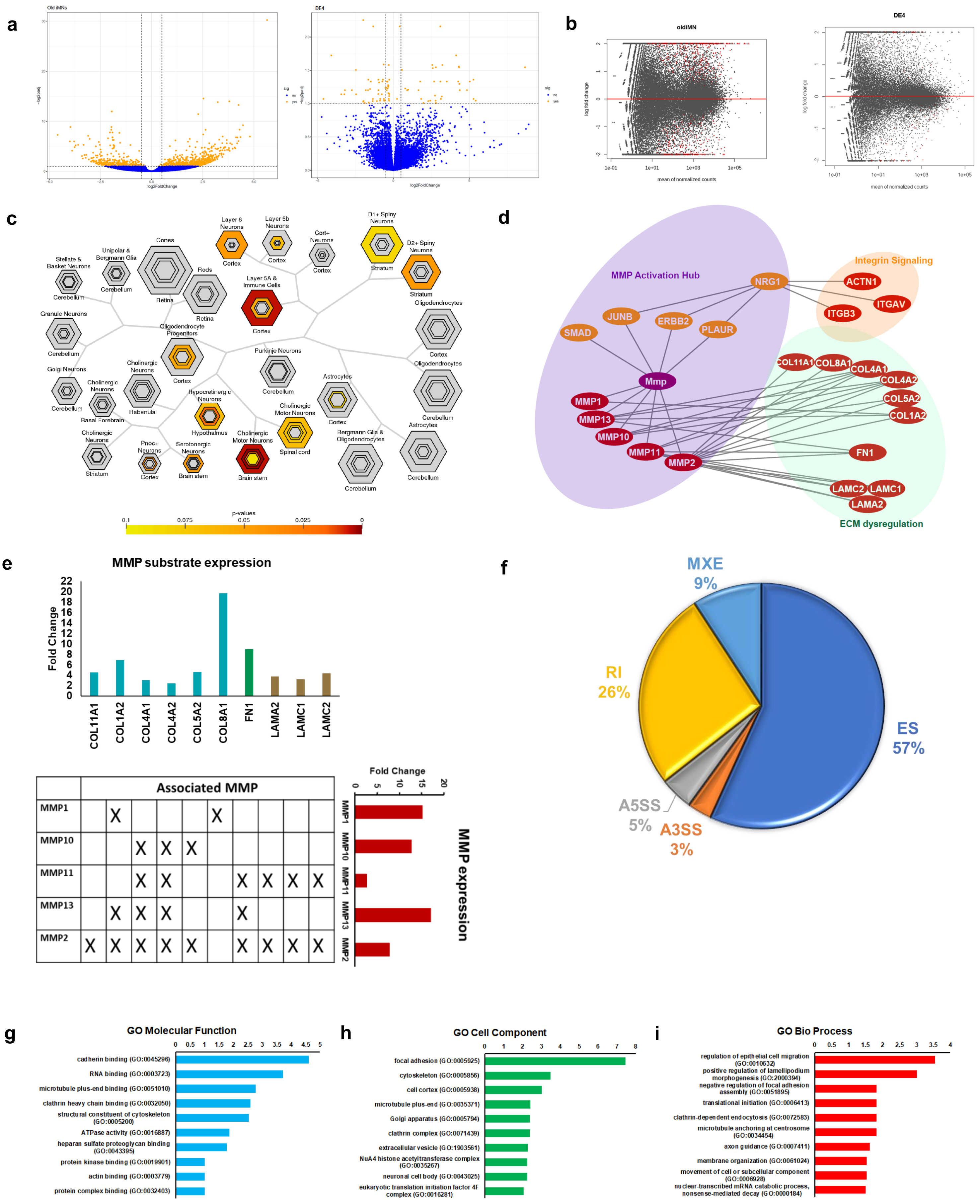

Supplementary Figure 7

a

| File Level Summary | MS Runs | Proteins | Peptides | Transitions |
|---|---|---|---|---|
| UniqueHits | 15 | 3844 | 23436 | 160686 |
| SharedHits | 15 | 5518 | 26102 | 179580 |

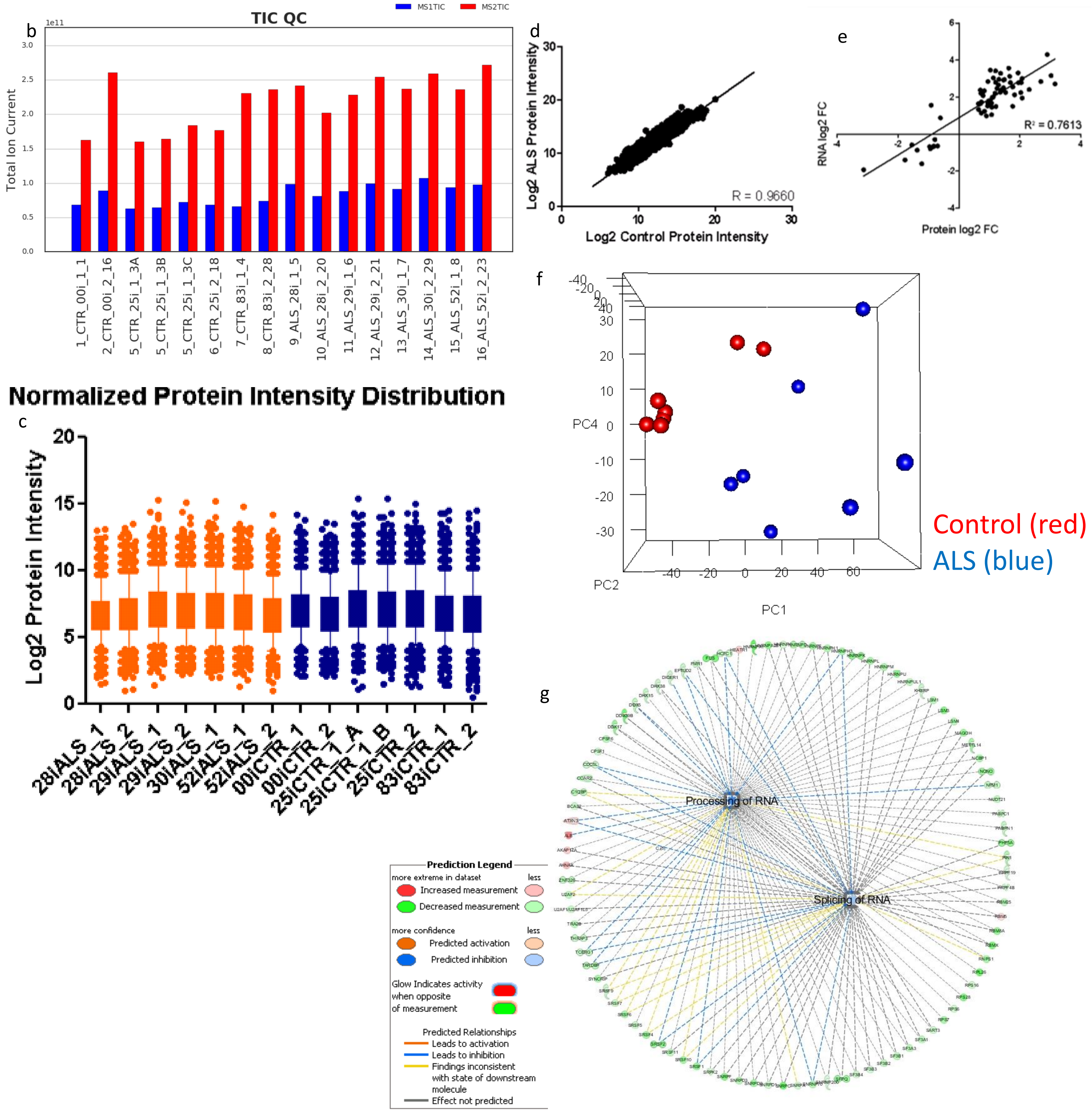

### Supplementary Figure 8

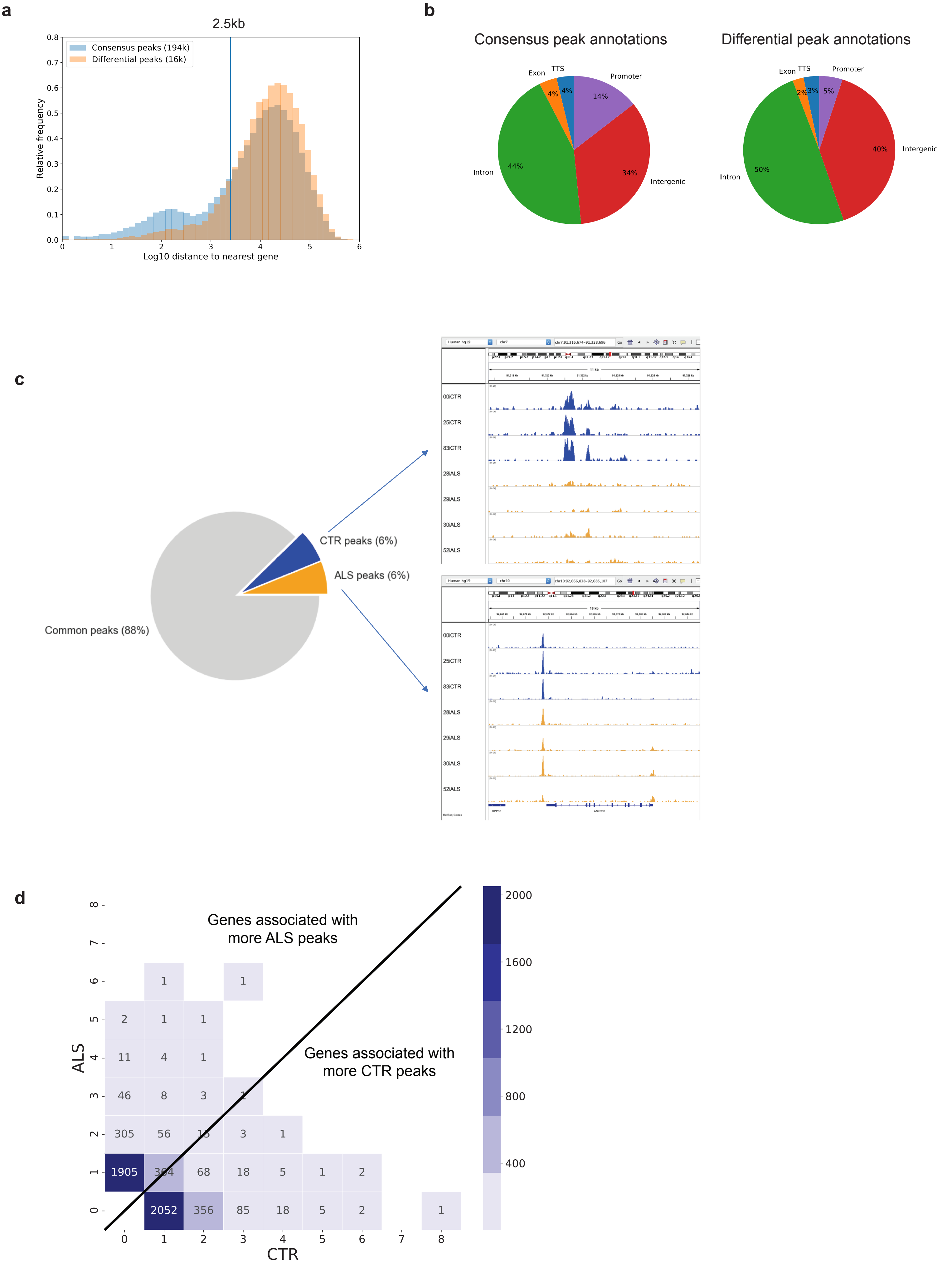

### Supplementary Figure 9

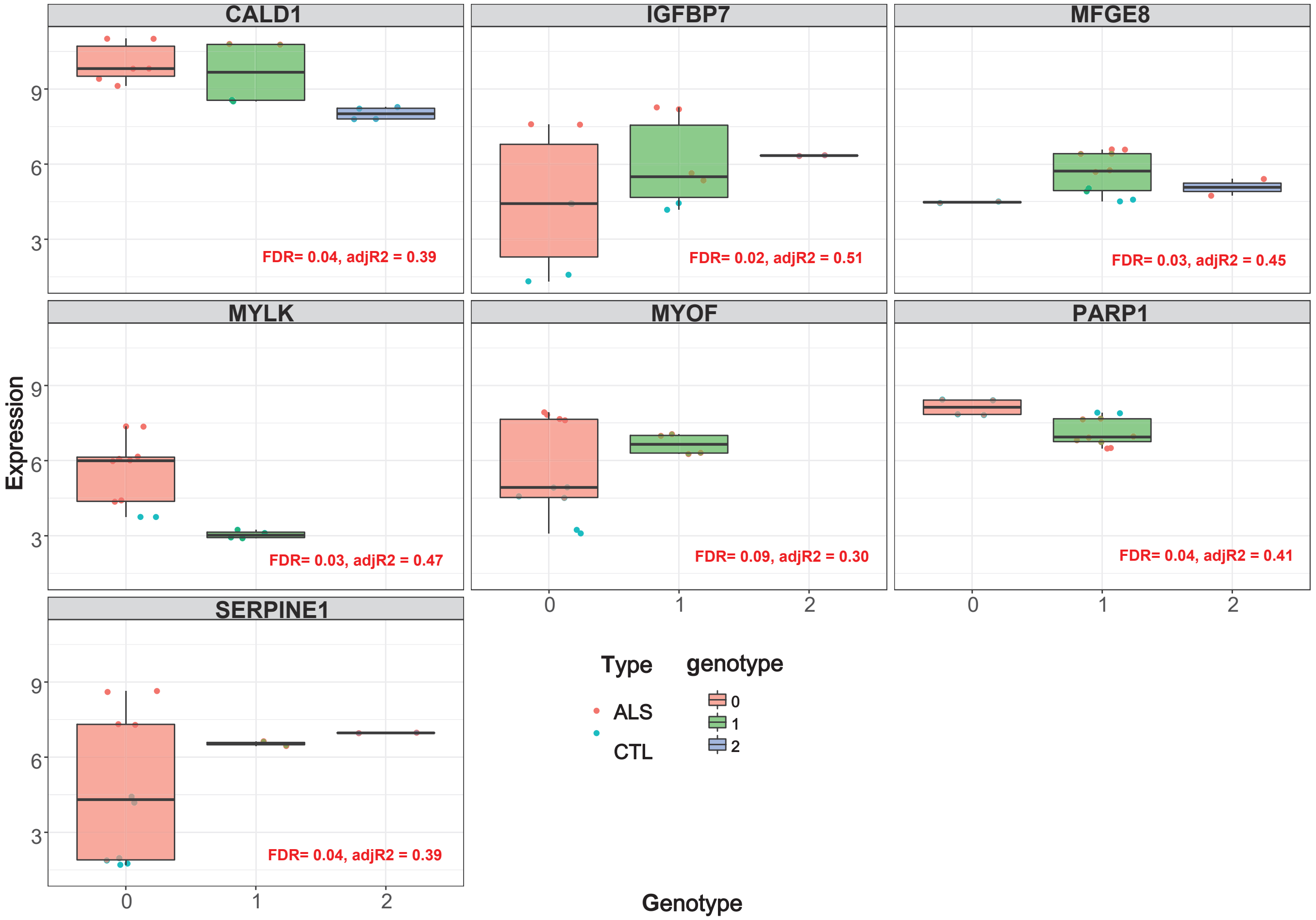

Supplementary Figure 10

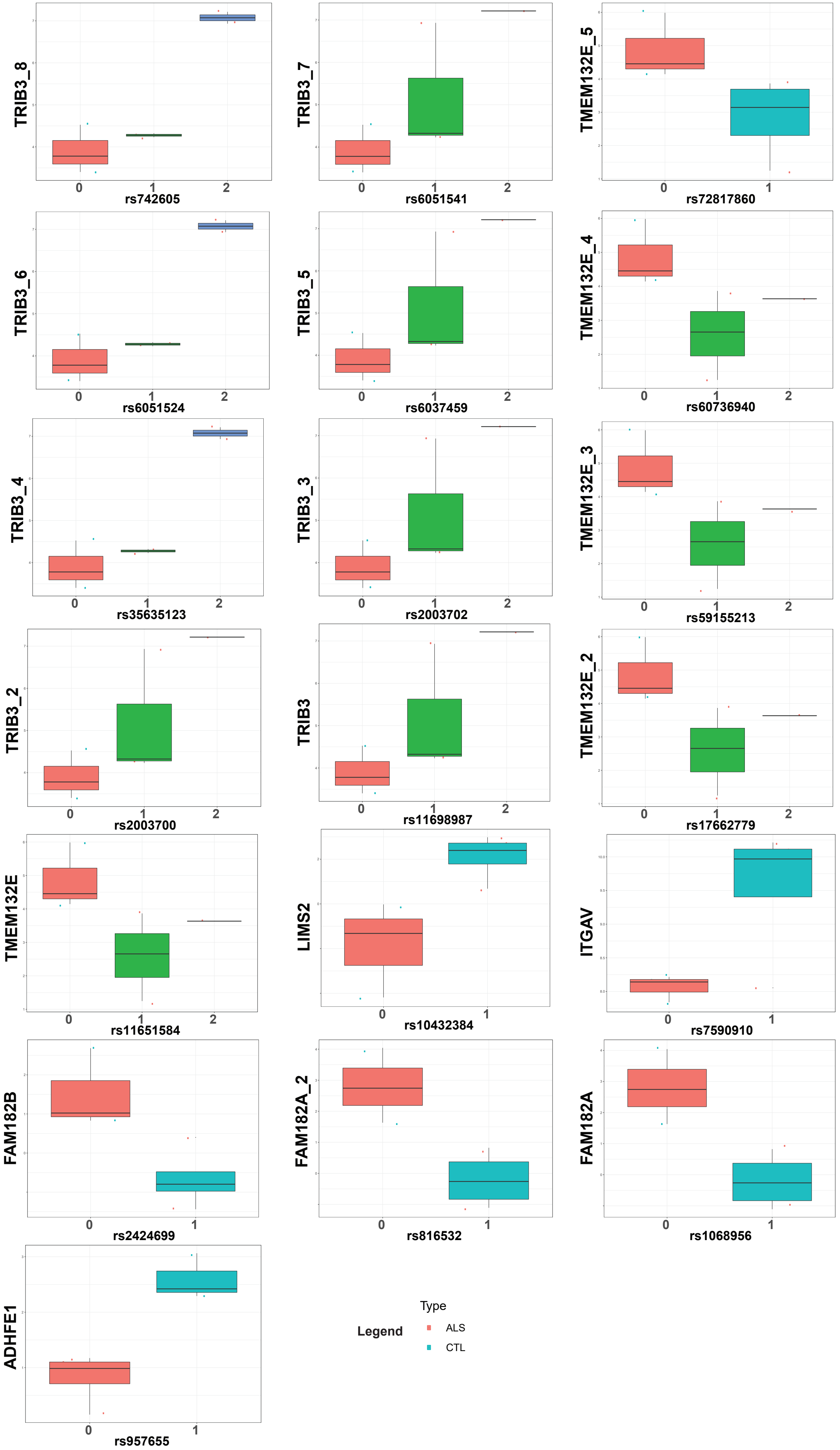

### Supplementary Figure 11

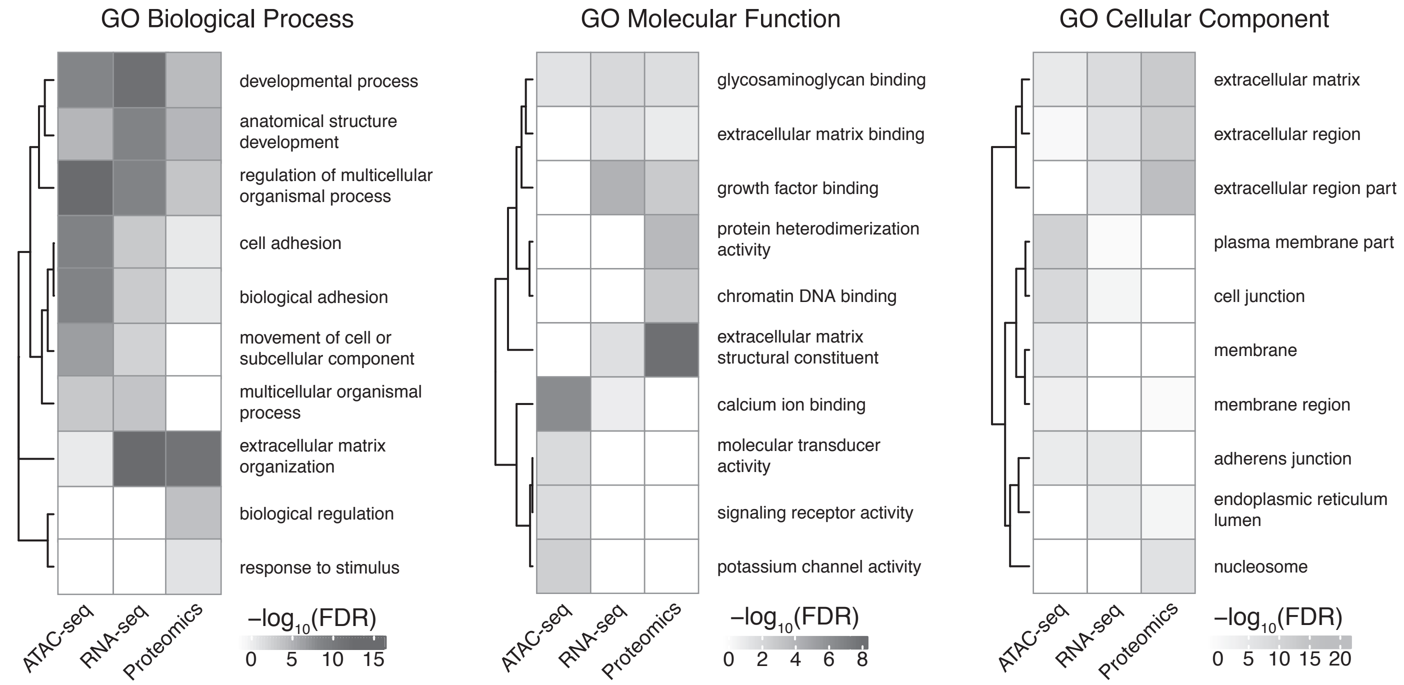

Supplementary Figure 12

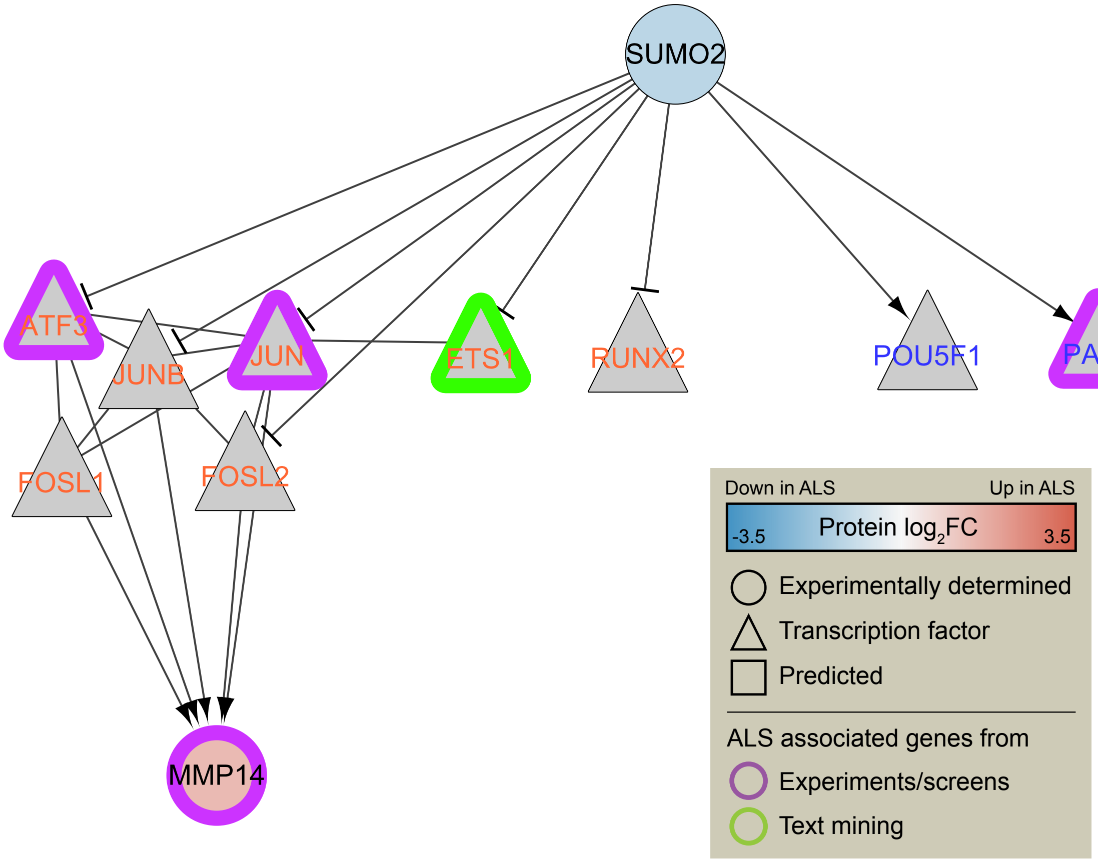

#### Supplementary Figure 13

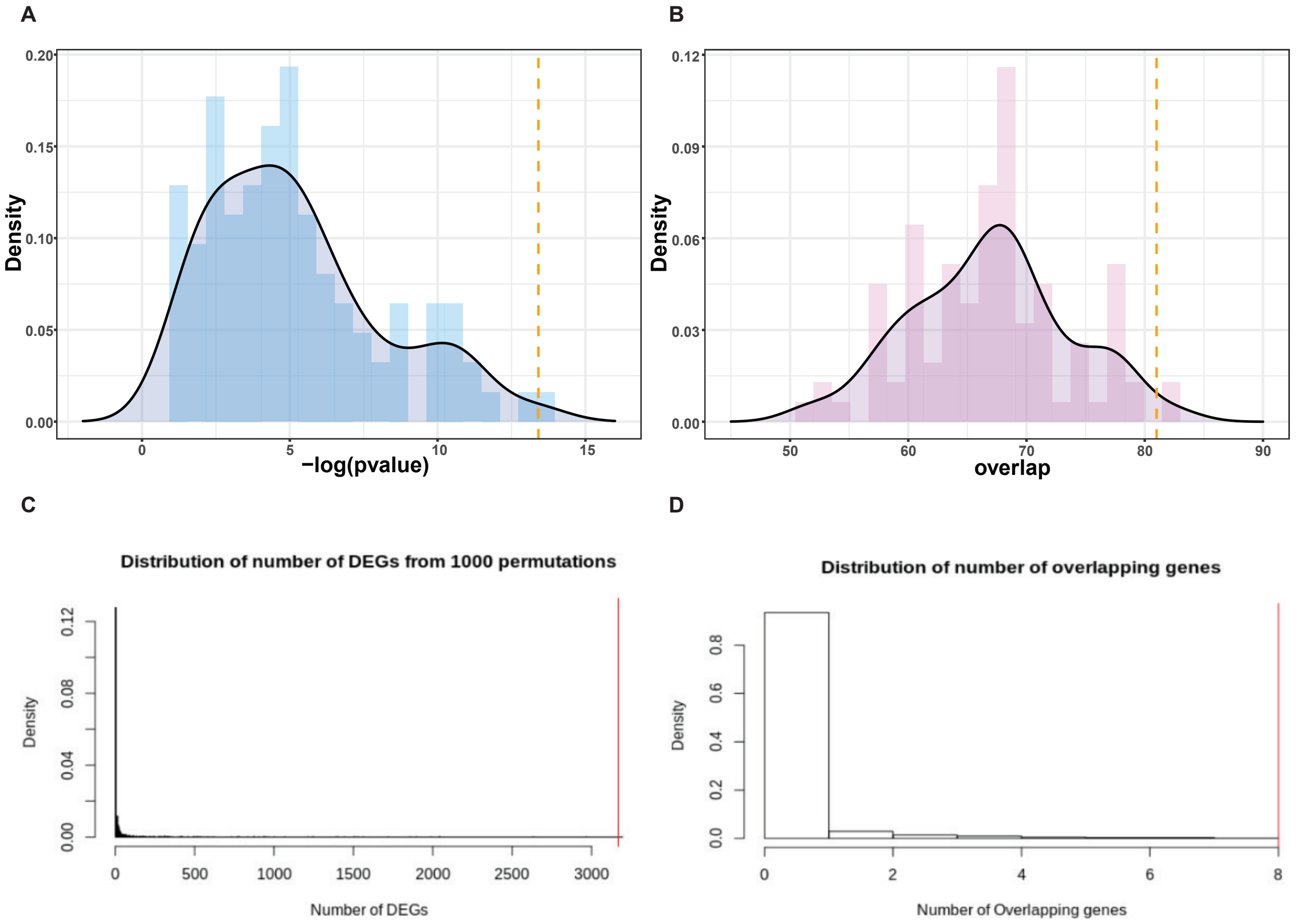

### Supplementary Figure 14

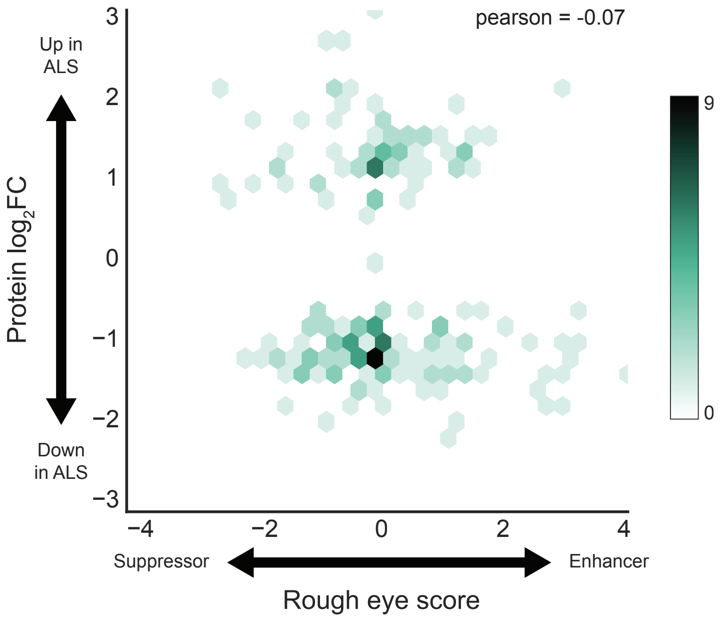

### Supplementary Figure 15

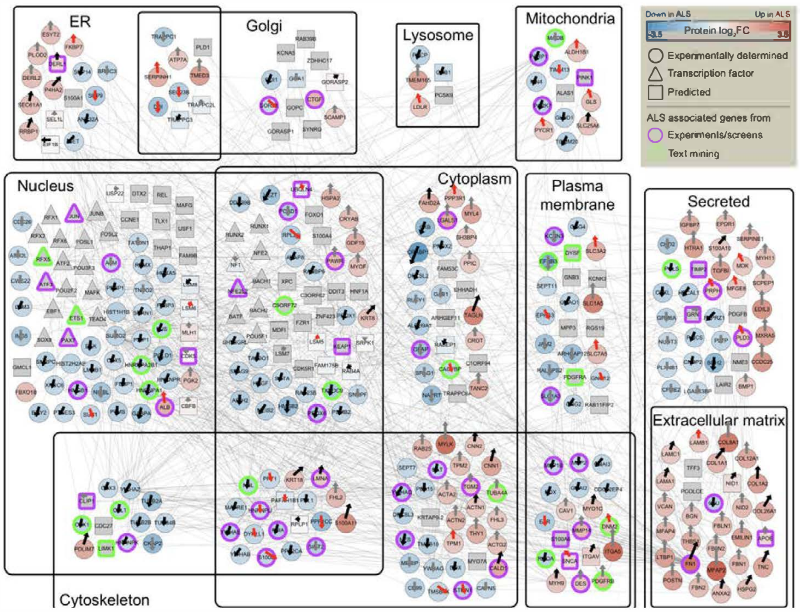

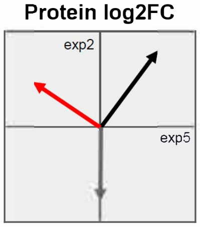

- **Black arrows** indicate nodes that are consistent, with log2FC in the same direction
- **Red arrows** indicate nodes that are inconsistent, with log2FC in opposite directions
- **Gray arrows** indicate nodes that are neither (not detected in both experiments)

- Null hypothesis: equal number of black and red arrows
- 133 black arrows
- 40 red arrows
- Significantly more black than red (pval=1.54E-12)
