## Supplemental tables for "An integrated multi-omic analysis of iPSC-derived motor neurons from C9ORF72 ALS patients"

Supplementary Table 1

| **Stage 1 (Day 0-Day 6)** | **Manufacturer** | **Catalog #** | **1X Concentration** |
|---|---|---|---|
| IMDM | Life Technologies | 12440061 | 47.5% |
| F12 | Life Technologies | 11765062 | 47.5% |
| NEAA | Life Technologies | 11140-50 | 1% |
| B27 | Life Technologies | 17504044 | 2% |
| N2 | Life Technologies | 17502048 | 1% |
| Anti/Anti | Life Technologies | 15240062 | 1% |
| LDN193189 | Cayman Chemical | 19396 | 0.2 µM |
| CHIR99021 | Xcess bioscience | M60002 | 3 µM |
| SB431542 | Cayman Chemical | 13031 | 10 µM |

Supplementary Table 2

| **Stage 2 Platedown (Day 6)** | **Manufacturer** | **Catalog #** | **1X Concentration** |
|---|---|---|---|
| IMDM | Life Technologies | 12440061 | 47.45% |
| F12 | Life Technologies | 11765062 | 47.45% |
| NEAA | Life Technologies | 11140-50 | 1% |
| B27 | Life Technologies | 17504044 | 2% |
| N2 | Life Technologies | 17502048 | 1% |
| Anti/Anti | Life Technologies | 15240062 | 1% |
| All-trans RA | Stemgent | 04-0021 | 0.1 µM |
| SAG | Cayman Chemical | 11914 | 1 µM |
| LDN193189 | Cayman Chemical | 19396 | 0.2 µM |
| CHIR99021 | Xcess bioscience | M60002 | 3 µM |
| SB431542 | Cayman Chemical | 13031 | 10 µM |
| Rock Inhibitor (Y-27632) | Stemcell Technologies | 72308 | 10 µM |

Supplementary Table 3

| **Stage 2 (Day 7-11)** | **Manufacturer** | **Catalog #** | **1X Concentration** |
|---|---|---|---|
| IMDM | Life Technologies | 12440061 | 47.5% |
| F12 | Life Technologies | 11765062 | 47.5% |
| NEAA | Life Technologies | 11140-50 | 1% |
| B27 | Life Technologies | 17504044 | 2% |
| N2 | Life Technologies | 17502048 | 1% |
| Anti/Anti | Life Technologies | 15240062 | 1% |
| All-trans RA | Stemgent | 04-0021 | 0.1 µM |
| SAG | Cayman Chemical | 11914 | 1 µM |
| LDN193189 | Cayman Chemical | 19396 | 0.2 µM |
| CHIR99021 | Xcess bioscience | M60002 | 3 µM |
| SB431542 | Cayman Chemical | 13031 | 10 µM |

Supplementary Table 4

| **Stage 3 (Day 12-Day18)** | **Manufacturer** | **Catalog #** | **1X Concentration** |
|---|---|---|---|
| IMDM | Life Technologies | 12440061 | 47.5% |
| F12 | Life Technologies | 11765062 | 47.5% |
| NEAA | Life Technologies | 11140-50 | 1% |
| B27 | Life Technologies | 17504044 | 2% |
| N2 | Life Technologies | 17502048 | 1% |
| Anti/Anti | Life Technologies | 15240062 | 1% |
| *SAG | Cayman Chemical | 11914 | 0.1 µM |
| db-cAMP | Millipore | 28745 | 0.1 µM |
| All-trans RA | Stemgent | 04-0021 | 0.5 µM |
| Compound E | Calbiochem | 565790 | 0.1 µM |
| DAPT | Cayman Chemical | 13197 | 2.5 µM |
| Ascorbic Acid | Sigma-Aldrich | A4403 | 200 ng/mL |
| BDNF (-80) | Peprotech | 450-02 | 10 ng/mL |
| GDNF (-80) | Peprotech | 450-10 | 10 ng/mL |

**Supplementary Table 5**

### C9ORF72-AMYOTROPHIC LATERAL SCLEROSIS

| iPSC Line | iPSC Method | Race/ Gender | Age of Onset | Age at Sampling | Mutation | *Source Tissue*/ Source | iPSC Source | Clinical remarks and family history |
|---|---|---|---|---|---|---|---|---|
| 28iALS-C9 | Episomal | W/M | 46 YR | 47 YR | *C9ORF72 exp.* ~800 $G_4C_2$ | *Fibroblast* WUSTL | CS iPSC Core | Father died of ALS age 70; sister died of ALS age 51; paternal cousins died age 50, 51 of ALS; brother with FTD. Site of onset: Left upper extremity |
| 29iALS-C9 | Episomal | W/M | 46 YR | 47 YR | *C9ORF72 exp.* ~800 $G_4C_2$ | *Fibroblast* WUSTL | CS iPSC Core | Maternal diagnosed with ALS; maternal grandfather diagnosed with dementia and parkinsonism. Site of onset: Left lower extremity |
| 30iALS-C9 | Episomal | W/F | 51 YR | 51 YR | *C9ORF72 exp.* ~70 $G_4C_2$ | *Fibroblast* WUSTL | CS iPSC Core | Mother with FTD maternal uncle and aunt, and two cousins died of ALS. Site of onset: Bulbar |
| 52iALS-C9 | Episomal | W/M | 45 YR | 48 YR | *C9ORF72 exp.* ~800 $G_4C_2$ | *Fibroblast* WUSTL | CS iPSC Core | Father died age 63 with dementia; paternal grandmother died of bulbar onset ALS. Site of onset: Left upper extremity |

### UNAFFECTED CONTROLS

| iPSC Line | iPSC Method | Race/ Gender | Age at Sampling | Diagnosis | Group/ Gene | *Source Tissue*/ Source | iPSC Source | Clinical remarks |
|---|---|---|---|---|---|---|---|---|
| 14iCTR | Episomal | W/F | ~35 YR | Clinically normal | SMA child's mother | *Fibroblast* Coriell | CS iPSC Core | 2 affected children with SMA; PCR analysis showed: donor subject has two copies of the SMN2; mother of 13iSMA. Apparently healthy non-fetal tissue |
| 00iCTR | Episomal | AA/M | 6 YR | Clinically normal | N/A | *Fibroblast* Coriell | CS iPSC Core | Apparently healthy non-fetal tissue |
| 83iCTR | Episomal | W/F | 21 YR | Clinically normal | HD negative | *Fibroblast* Coriell | CS iPSC Core | Asymptomatic; HD Gene-negative, Apparently healthy non-fetal tissue |
| 25iCTR | Episomal | W/M | 76 YR | Clinically normal | HD negative | *Fibroblast* Coriell | CS iPSC Core | Asymptomatic; HD Gene-negative. Apparently healthy non-fetal tissue |

**Table S5**: Clinical and reprogramming details for the ALS-C9 and control iPSC lines used in the NeuroLINCS study. W: White; AA: African American; M: Male; F: Female; CS: Cedars-Sinai; HD: Huntington Disease.

**Supplementary Table 6**

| Variant Type | Total Variants |
|---|---|
| Total variants in the 3 control iPSC lines | 9,197,462 |
| Exonic functional | 57,910 |
| Rare Exonic functional < 1% frequent or novel (no frequency information) | 12,898 |
| Regulatory | 4,947,479 |
| Total variants in iPSC lines with C9ORF72 mutation | 8,818,235 |
| Exonic functional | 55,815 |
| Rare Exonic functional < 1% frequent or novel (no frequency information) | 8,225 |
| Regulatory C9ORF72 | 4,740,665 |

**Table S6:** Summary Table of all variants in iPSC lines from healthy volunteers and people with ALS due to C9ORF72 mutation. The total number of variants that are exonic functional are reported. These are nonsynonymous variants which include missense, splicing. frameshift, non-frameshift, stop-gain and start loss variants only. For regulatory variants, we filtered for variants that are in intergenic and regulatory regions. We report the variant as found next to the closest gene, these will be either in the 5’ or 3’ UTR, intronic, upstream and downstream up to 4 KBs from the start and stop of a gene.

**Supplementary Table 7**

| Specific ALS Associated Variants Found (ALSoD database) | Control Line | C9ORF72 Line |
|---|---|---|
| OPTN c.293T>A:p.M98K:(16Exons):exon5:missense (3.0% frequent) | CS83iCTR-33n1 | CS28iALS-C9n2 |
| ALS2 c.280A>G:p.I94V(4Exons):exon4:missense (2.1% frequent) | - | CS52iALS-C9n6 |
| DIAPH3 c.974C>T:p.P325L:(16Exons):exon10:missense (3.6 % frequent) | CS83iCTR-33n1 | - |

**Table S7:** Summary Table of all ALS variants in the control and C9ORF72 lines. ALS specific variants were found in one of the CS83iCTR-33n1 and 2 of the C9ORF72 lines.

**Supplementary Table 8**

| Variants that shared across _ and are in _ | Total Variants | Variants that overlap with RNA Seq data | Variants that overlap with ATAC Seq data | Non-synonymous (includes Start-loss (STL) | Frame Shift (FS) | Splicing (S) | Stop-gain (SG) | Stop-loss (SPL) |
|---|---|---|---|---|---|---|---|---|
| 3 controls and all C9ORF72 (cases) lines | 98,897 | 4,908 | 847 | 29,052 | 736 | 872 | 365 | 43 |
| 4 cases, 0 controls | 206 | 16 | 1 | 82 | 2 | 1 | 0 | 0 |
| 4 cases, 1 control | 910 | 62 | 8 | 312 | 4 | 11 | 8 | 1 |
| 4 cases, 2 controls | 2,774 | 186 | 22 | 1,047 | 16 | 37 | 8 | 2 |
| 3 cases, 0 controls | 799 | 49 | 7 | 290 | 9 | 11 | 4 | 1 |
| 3 cases, 1 control | 2,259 | 155 | 33 | 828 | 11 | 18 | 5 | 2 |
| 2 cases, 0 controls | 2,784 | 195 | 37 | 1,021 | 37 | 17 | 17 | 0 |
| 1 case, 3 controls | 638 | 52 | 4 | 222 | 4 | 11 | 0 | 0 |
| 0 case, 3 controls | 218 | 7 | 4 | 96 | 1 | 2 | 0 | 1 |
| 0 case, 2 controls | 1,405 | 79 | 13 | 509 | 11 | 20 | 10 | 3 |
| Total enriched in cases or controls | 11,993 | 801 | 129 | 4,407 | 95 | 128 | 52 | 10 |

**Table S8:** Summary table of the exonic functional variants within all lines the variants that are enriched in either the cases or controls.

**Table S9:** List of genes analyzed for statistically significant differences from RNAseq. See Excel sheet Table S6.

**Table S10**: Summary of Proteomics. See Excel Sheet Table S7 and corresponding tabs.

**Table S11:** Summary table of the exonic functional variants within all lines the variants that are enriched in either the cases or controls.

**Table S12**: Genes used for linear regression analysis of gene variant and expression comparisons.

**Table S13**: List of ALS associated genes from text mining sources[1] and experimental sources[2,3].

**Table S14:** Summary table of RNAi lines used and full G4C2 fly screen results.

**Table S15:** Proteomics fold changes (ALS/CTR) and false discovery rates (FDR) from the original and validation experiments for proteins in the fly network (Figure 6).

1. Koscielny, G., *et al.* Open Targets: a platform for therapeutic target identification and validation. *Nucleic acids research* **45**, D985-D994 (2017)5210543.
2. Freibaum, B.D., *et al.* GGGGCC repeat expansion in C9orf72 compromises nucleocytoplasmic transport. *Nature* **525**, 129-133 (2015)4631399.
3. Jovicic, A., *et al.* Modifiers of C9orf72 dipeptide repeat toxicity connect nucleocytoplasmic transport defects to FTD/ALS. *Nature neuroscience* **18**, 1226-1229 (2015)4552077.
